# Single-nucleus RNA-seq identifies divergent populations of FSHD2 myotube nuclei

**DOI:** 10.1101/478636

**Authors:** Shan Jiang, Katherine Williams, Xiangduo Kong, Weihua Zeng, Xinyi Ma, Rabi Tawil, Kyoko Yokomori, Ali Mortazavi

## Abstract

FSHD is characterized by the misexpression of *DUX4* in skeletal muscle. However, *DUX4* is lowly expressed in patient samples and analysis of the consequences of *DUX4* expression has largely relied on artificial overexpression. To better understand the native expression profile of *DUX4* and its targets, we performed pooled RNA-seq differentiation time-course in FSHD2 patient-derived primary myoblasts and identified early-and late-induced sets of FSHD-associated genes. Using single-cell and single-nucleus RNA-seq on FSHD2 myoblasts and myotubes respectively, we captured *DUX4* expression in single-nuclei and found that only some *DUX4* targets are coexpressed. We identified two populations of FSHD myotube nuclei with distinct transcriptional profiles. One population is highly enriched with *DUX4* and FSHD related genes, including the *DUX4* paralog *DUXA* (“FSHD-Hi”). The other population has no expression of *DUX4* and expresses low amounts of FSHD related genes (“FSHD-Lo”), but is marked by the expression of *CYTL1* and *CHI3L1*. “FSHD-Hi” myotube nuclei upregulated a set of transcription factors (TFs) that may form a self-sustaining network of gene dysregulation, which perpetuates this disease after *DUX4* is no longer expressed.

## Introduction

Fascioscapulohumeral muscular dystrophy (FSHD) is one of the most common inherited muscular dystrophies and is characterized by progressive wasting of facial, shoulder and upper arm musculature [1]. The most common form of FSHD, FSHD1 (>95% of cases), is linked to the mono-allelic contraction of the D4Z4 macrosatellite repeat array on chromosome 4q from 11-100 units to 1-10 units, with each 3.3 kb repeat containing the open reading frame for the double-homeobox transcription factor DUX4 [2-4]. In contrast, FSHD2 (<5% of FSHD cases) has no contraction of the chromosome 4q repeat array. More than 80% of FSHD2 cases are characterized by recurring mutations in the chromatin modifier SMCHD1 (Structural Maintenance of Chromosomes flexible Hinge Domain-containing protein 1) on chromosome 18 [5]. SMCHD1 is important for maintenance of DNA methylation and epigenetic silencing of multiple genomic loci, including the D4Z4 repeat array [5]. Studies have also found that SMCHD1 mutations can act as disease modifiers in severe cases of FSHD1 [6, 7].

FSHD is associated with the expression of the full-length *DUX4* transcript (*DUX4fl*) which is stabilized by a specific single-nucleotide polymorphism in the chromosomal region distal to the last D4Z4 repeat by creating a canonical polyadenylation signal [8-10]. *DUX4fl* encodes a transcriptional activator with a double-homeobox domain that binds to a specific sequence motif upstream of its target genes in the genome [3, 4]. Normally, *DUX4* is briefly expressed in 4-cell human embryos when it activates genes for zygote genome activation (ZGA) [11-13]. In muscle cells, overexpression of *DUX4fl* causes differentiation defects and cytotoxicity in human and mouse myoblasts [14, 15]. However, the endogenous *DUX4fl* is expressed at extremely low levels in FSHD and DUX4 protein is only detected in 0.001% and 0.005% of patient myoblasts and myotubes, respectively, *in vitro* [16]. The relationship of DUX4-positive and -negative cells and whether DUX4-negative patient cells contribute to the disease is unclear. The regulation of *DUX4* expression is controlled by multiple epigenetic processes. D4Z4 repeats are normally heterochromatic with DNA hypermethylation and histone H3 lysine 9 trimethylation (H3K9me3), which are lost in FSHD1 and FSHD2 [17, 18]. The depletion of SMCHD1, which binds to D4Z4 repeats in an H3K9me3-dependent fashion [2], results in *DUX4fl* upregulation and mutations throughout the gene correlate with CpG hypomethylation in D4Z4 repeats [19].

Here we focused on the SMCHD1-mutated FSHD2 subtype in order to characterize the heterogeneity of *DUX4* and FSHD-induced target gene expression at the single-cell level using *in vitro* differentiation of primary FSHD2 patient-derived myoblasts into myotubes. Although FSHD2 represents a minor population of FSHD cases, patient cells exhibit comparable clinical and gene expression phenotype as FSHD1 [20]. We used two FSHD2 patient samples with defined genetic mutations of *SMCHD1* and significant DNA hypomethylation of D4Z4. Using pooled RNA-seq, we profiled gene expression patterns during a differentiation time-course and identified candidate disease-related key genes (i.e. FSHD-induced genes) that are upregulated specifically in FSHD cells by comparing expression profiles between FSHD2 and control. We found that about 40% of differentially expressed genes are known FSHD-associated genes while the others may be potential new candidates involved in the progression of the disease. We then used single-cell RNA-seq in myoblasts and single-nucleus RNA-seq [21] in 3-day post-differentiation myotubes to characterize the expression patterns of *DUX4* and other FSHD-induced genes. We sucessfully detected the first set of single nuclei with *DUX4* expression (*DUX4*-detected) from FSHD myotubes and found that they do not express all the FSHD-induced genes whereas a much larger set of FSHD myotube nuclei express FSHD-induced genes. Using pseudotime analysis on the single-cell/single-nucleus data, we also found two interesting bifurcation time points that (1) separate FSHD2 myotube nuclei from control myotube nuclei and (2) distinguish FSHD myotube nuclei that have low expression of FSHD-induced genes (termed “FSHD-Lo”) from the nuclei with high expression of FSHD-induced genes (termed “FSHD-Hi”) nuclei. FSHD-Hi nuclei, either with or without *DUX4* expression, show the highest enrichment of FSHD differentialy expressed genes observed in the population-based analysis. We found that *DUX4* and its paralogs, *DUXA* and *DUXB*, are expressed in “FSHD-Hi” and “FSHD-Lo” nuclei respectively and have distinct coexpression patterns with other FSHD-induced genes. These two populations of FSHD myotube nuclei may enter distinct biological programs driven by differential expression of specific sets of transcrition factors downstream of DUX4.

## Results

### Upregulation of FSHD-induced genes during FSHD2 myotube differentiation

Previous studies indicated that *DUX4* is upregulated during FSHD patient myoblast differentiation [22]. In order to understand the temporal expression differences between FSHD2 patient-derived and control myoblasts, we differentiated these *in vitro* to measure the dynamics of gene expression in a 6-day time-course using conventional pooled RNA sequencing (RNA-seq) (Figure 1A, S1) (Methods). We used two independent primary control myoblast samples, Control-1 and Control-2, and two independent primary FSHD2 myoblast samples, FSHD2-1 and FSHD2-2, which have known *SMCHD1* mutations. In addition, we compared our time-course to the differentiation of the immortalized human myoblast cell line KD3 [23] which was used previously to study the fate heterogeneity during myotube differentiation by single-nucleus RNA-seq [21]. After sequencing two biological replicate RNA samples for each of the five cell populations every day for six days, we filtered out lowly expressed genes across all 60 samples and performed principal component analysis (PCA) with 10,767 genes (Figure 1B). We observed that the days of differentiation aligned to each other across cell lines following a clear trajectory of myogenesis (PC1, 34.3% variance in expression). FSHD2 and control clearly separated from KD3 by the second principal component (PC2, 19.1% variance in expression) which may be due to the immortalization of KD3 and/or the different muscles from which they were derived (control and FSHD2-2 myoblast cells were from quadriceps, and FSHD2-1 myoblast cells were from tibia; KD3 from trunk muscle) (Figure S2). We also found that the two FSHD2 cell lines had similar variance in expression to the two control cell lines at the early differentiation timepoints but separated distinctly for days 3 to 5 in two principal components with known genes upregulated in FSHD driving the variance (PC4, 6.6% variance in expression; PC7, 3.6% variance in expression) (Figure 1B, Table S1). Thus, FSHD2 patient-derived myotubes can be distinguished from control and KD3 cells by day 3 of differentiation when profiling transcriptomes at the population level.

**Figure 1.**
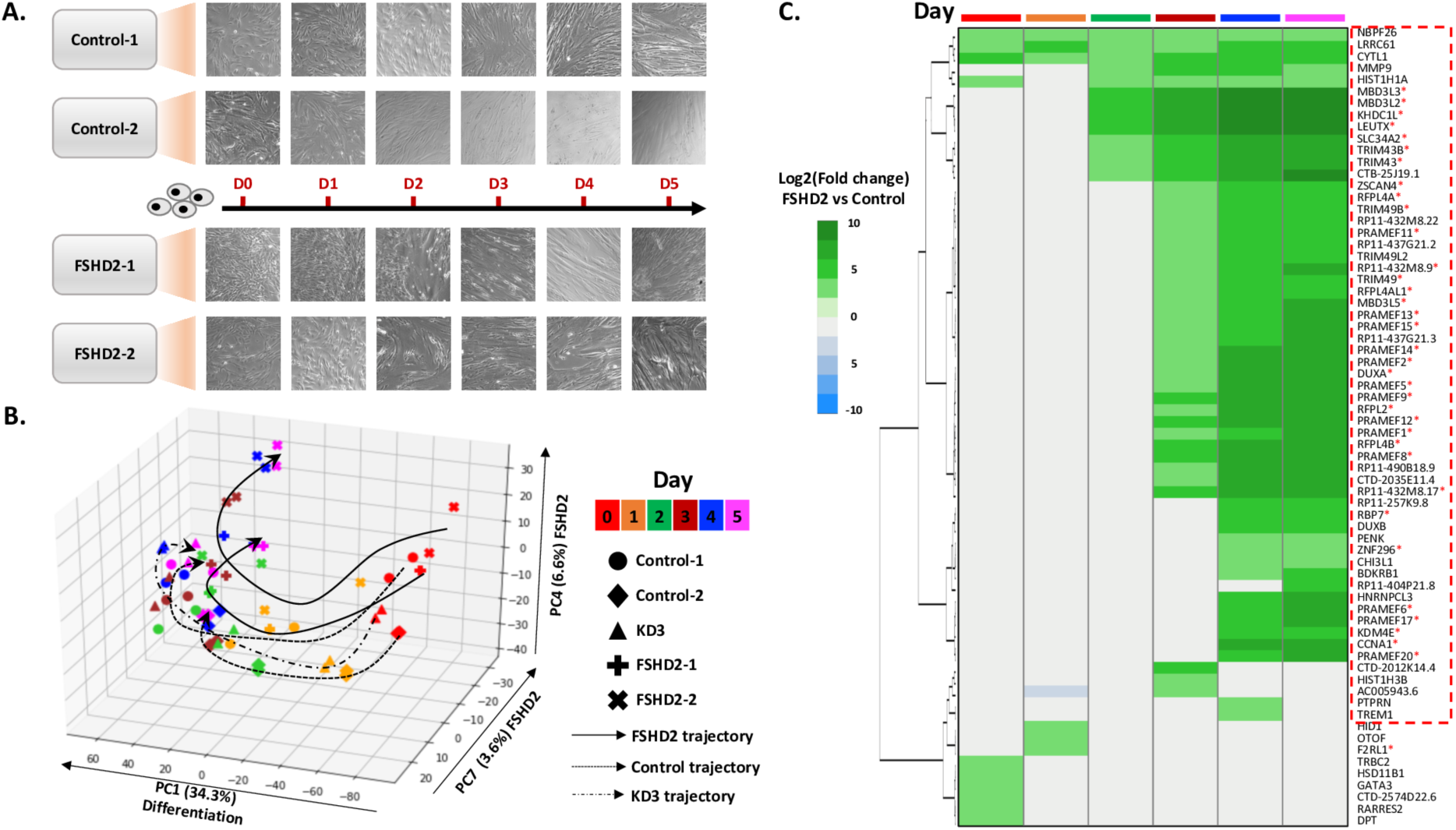
Upregulation of FSHD-induced genes starting at day 2 identified in pooled RNA-seq time-course. **(A)** Differentiation time-course of control and FSHD2 patient-derived myoblast to myotube. Morphology changes are shown per day during differentiation. **(B)** Principal component analysis (PCA) on control, FSHD2 and KD3 myoblast differentiation time-course. Gene expression level is measured each day for duplicates by using RNA-seq. Cell types are labeled by shape, and time-points are labeled by color. **(C)** Hierarchical heatmap of expression fold changes (FSHD2 vs control) on up-regulated genes in FSHD2 per day. A total of 59 known and potential DUX4 targets that are up-regulated at day 3 or later stage are labeled out with dashed red box. Genes that have been published to be induced by DUX4 are labeled with a red star.

In order to comprehensively characterize the expression of *DUX4* and its associated upregulated genes during myogenesis, we examined differentially expressed genes (log2|FC|>3) by comparing between FSHD2 and control at each day of the time-course (Methods). We identified 103 differentially expressed genes (68 upregulated, 35 downregulated) in FSHD2 across the time-course (Figure 1C, Table S2). Of the 68 upregulated genes, we call 36 as confirmed DUX4 targets since they have been shown to be overexpressed in DUX4-positive myotubes [22] or in FSHD muscle biopsies, and/or they have DUX4 binding sites confirmed by ChIP-seq [20] (Figure 1C red astericks, S3). We define 59 “FSHD-induced genes” as the set of known DUX4 target genes, genes upregulated in previous FSHD studies, and novel genes upregulated across this FSHD2 differentiation time-course (Figure 1C red box). Previously published FSHD-induced genes [20, 22, 24] were upregulated in waves starting at day 2, such as *TRIM43*, *MBD3L2*, *MBD3L3*, *KHDC1L* and *LEUTX*, followed by day 3, such as *PRAMEF1*, *PRAMEF2*, *PRAMEF12*, *ZSCAN4* and *MBD3L5*, and day 4, such as *CCNA1*, *KDM4E*, *PRAMEF17*, *PRAMEF6* [20] (Figure 1C, S3). After being significantly upregulated, most FSHD-induced genes remained upregulated through the end of the time-course (Figure 1C). As DUX4 protein was shown to be transiently expressed [22], the continued up-regulation of these genes could be due to a self-supporting network of gene regulation initially induced by DUX4 then possibly mediated by transcription factors and chromatin remodelers, such as ZSCAN4 and MBD3L2. Interetingly, *DUXA* and *DUXB*, which are paralogs of *DUX4* [25], were also stably activated starting at days 3 and 4, respectively, in FSHD2 samples suggesting their potential contribution in FSHD-related pathways.

In order to identify temporal patterns of expression, we used maSigPro [26] to cluster the differentially expressed genes into four clusters with genes upregulated in FSHD2 separating into early-(clusters 1) and late-induced expression patterns (cluster 4) (Figure 2A, 2B, 2D) (Methods). Importantly, known DUX4-associated upregulated genes [20] represent over 70% of the genes induced in later stage, indicating that other genes within the same clusters could be novel candidate target genes for DUX4 in FSHD2. Genes induced at the later stage (cluster 4) were highly enriched in GO terms for negative regulation of cell differentiation (p = 1 × 10^−9.76^) and methylation-dependent chromatin silencing (p = 1 × 10^−5.56^) (Figure 2D, 2E, Table S3) while genes induced at the beginning of differentiation (cluster 1) were enriched in GO terms for leukocyte migration (p = 1 × 10^−5.24^), humoral immune response (p = 1 × 10^−2.96^), response to oxidative stress (p = 1 × 10^−2.67^) and chromatin organization (p = 1 × 10^−1.96^) (Figure 2B, 2C, Table S3). The other two clusters represent genes upregulated in both FSHD2 and control differentiation (cluster 2) or only in Control-2 (cluster 3) (Figure S4, Table S3). In summary, we found a set of genes significantly upregulated in differentiating FSHD2 myotubes with distinct early and late dynamics with most of the characteristic FSHD-induced genes activated by day 3.

**Figure 2.**
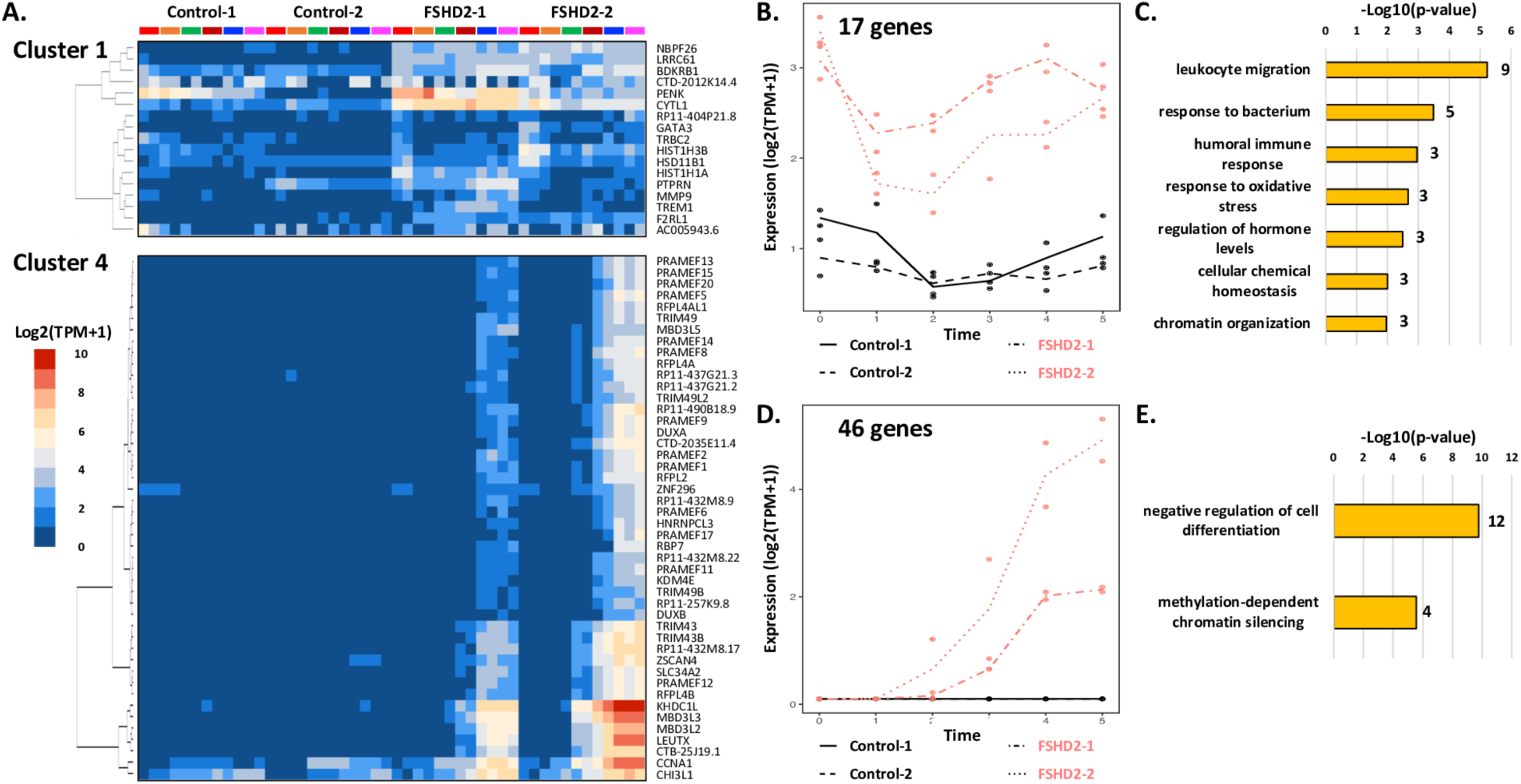
Early-and late-induced expression patterns of FSHD-induced genes. **(A)** 103 differentially expressed genes clustered into four clusters by maSigPro based on their expression patterns during control and FSHD2 differentiation time-course. Gene expression heatmap of two of the clusters of genes that are upregulated in FSHD2. Genes in cluster 1 are upregulated at early stage of differentiation while genes in cluster 4 are upregulated at the later stage of differentiation. **(B)** Expression profiles of the 17 genes in cluster 1 across four cell types during differentiation. **(C)** Significant gene ontology terms associated with the 17 genes in cluster 1 (FDR < 0.05). **(D)** Expression profiles of the 46 genes in cluster 4 across four cell types during differentiation. **(E)** Significant gene ontology terms associated with the 46 genes in cluster 4 (FDR < 0.05).

### Detection of nuclei with *DUX4* expression from FSHD2 myotubes using single-nucleus RNA-seq

Although we failed to detect *DUX4* in our pooled RNA-seq, the upregulation of FSHD-induced genes was nevertheless observed during myotube differentiation specifically in FSHD2 samples. We wondered whether the expression of FSHD-induced genes is seen in every cell and whether the expression of *DUX4* and DUX4-target genes were indeed present only in a subset of cells. We therefore performed single-cell RNA-seq on undifferentiated myoblasts and single-nucleus RNA-seq on myotubes using the Fluidigm C1 platform [21] at day 3 of differentiation using control and FSHD2 primary cells (Figure S5A). Day 3 was chosen as it was the first day of robust FSHD-induced gene expression in the differentiation time-course thereby allowing us to observe early transcriptional changes. Given the unique profile of Control-2 (Figure S4D), we used Control-1 for single-cell/single-nucleus experiments. Additionally, we selected FSHD2-2 based on the higher expression level of FSHD-induced genes compared to FSHD2-1 during differentiation (Figure 2A, D). We also compared these results to our previously published data on single-cell/single-nucleus RNA-seq in immortalized KD3 cells [21]. As quality control that our single cell data matched our pooled time-course, we first pooled reads from all single cells/single nuclei for each cell type and performed incremental PCA with the pooled time-course RNA-seq samples (Figure S5B, S5C). As expected, the pooled single-cell myoblasts clustered with day 0 samples in both control and FSHD2. For the pooled myotube single nuclei, FSHD2 replicate 1 (FSHD2 R1) aligned with day 3 of the FSHD2 time-course, but FSHD2 replicate 2 (FSHD2 R2) located between control and FSHD2 day 3 in the time-course (Figure S5C). This suggests variable differentiation efficiencies for the two replicates, which could be caused by subtle differences in seeding density.

Importantly, we found that 3 out of 79 (3.8%) nuclei in FSHD2 R1 showed high expression of *DUX4* (11.24 TPM, 34.15 TPM and 68.49 TPM) while we detected no *DUX4*-detected nuclei in FSHD2 R2, revealing the high level of heterogeneity in the FSHD2 cell population with *DUX4* only expressed in a small fraction of nuclei. We then analyzed the global profiles of the single-cell transcriptomes using PCA analysis and found that all 3 *DUX4*-detected nuclei as well as other FSHD2 R1 nuclei clearly separated from FSHD2 R2 and control myotube nuclei (Figure 3A). We analyzed 59 FSHD-induced genes specifically upregulated at day 3 or later during our pooled time-course of FSHD2 differentiation (Figure 1C) and observed that these genes showed significant enrichment in FSHD2 R1 myotube nuclei compared with control myotube nuclei (p < 2.2 × 10^−16^). Nuclei with the highest enrichment clustered with the 3 *DUX4*-detected nuclei, and thus we labeled this group of nuclei “FSHD-induced genes high” (“FSHD-Hi”) (Figure 3B, S6A). The FSHD2 R2 myotube nuclei also showed significantly higher enrichment of FSHD-induced genes than control myotube nuclei (p < 2.2 × 10^−16^) but had fewer FSHD-induced genes expressed than the “FSHD-Hi” group, and therefore this group of nuclei was labeled “FSHD-induced genes low” (“FSHD-Lo”) (Figure 3B, S6A). However, genes downregulated during FSHD2 myogenesis were comparable across all types of cells/nuclei but with significantly lower detection in the “FSHD-Hi” nuclei (Figure S6A, S6B). In summary, we detected three different patient myotube nuclei populations: (1) a set of 3 nuclei that express endogenous *DUX4* (*DUX4*-detected) and FSHD-induced genes (“FSHD-Hi”); (2) a larger set of 76 nuclei that express FSHD-induced genes but no detectable *DUX4* (“FSHD-Hi”); and (3) 60 nuclei that are clearly different from control nuclei but nevertheless with no *DUX4* and significantly lower FSHD-induced gene expression (“FSHD-Lo”).

**Figure 3.**
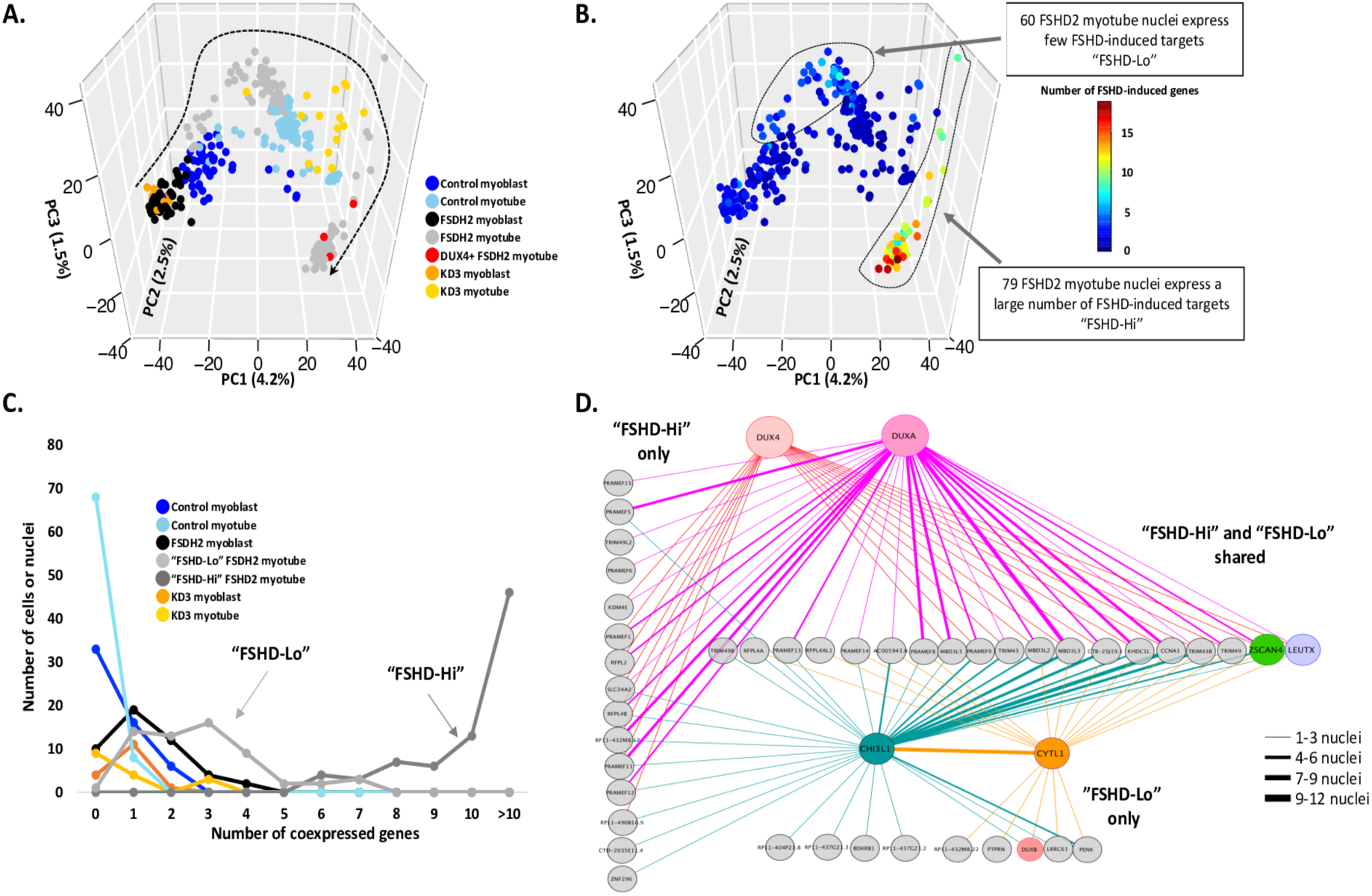
FSHD2 myotube nuclei can be separated into two clusters with differential expression of FSHD-induced genes. **(A)** PCA of single-cell (for myoblast) and single-nucleus (for myotube) RNA-seq data for control, FSHD2 and KD3. Cell types are labeled by distinct colored points and 3 DUX4-detected FSHD2 myotube nuclei are specifically labeled in red. **(B)** PCA from panel (A) colored by the number of FSHD-induced genes in FSHD2 that are up-regulated at day 3 or later stages. **(C)** Summary of the number of co-expressed genes (47 of the FSHD-induced genes that show continuous up-regulation from day 2) in different cell types. Cell types are labeled by color. **(D)** Coexpression network between DUX4, DUXA, CHI3L1, CYTL1 and other FSHD-induced genes in “FSHD-Hi” and “FSHD-Lo” nuclei.

To assess whether these groups of nuclei have distinct expression of FSHD-induced genes, we determined the coexpression patterns between a subset of FSHD-induced genes which had variable expression in the single cells and nuclei (excluding 7 genes *MMP9*, *TREM1*, *HNRNPCL3*, *PRAMEF17*, *PRAMEF2*, *PRAMEF20* and *RBP7* that were not detected as well as *NBPF26* that was expressed in all single cells and nuclei). We found that all myoblast cells and control or KD3 myotube nuclei typically express rarely more than 2 FSHD-induced genes (Figure 3C), whereas “FSHD-Lo” nuclei coexpress between 1 to 5 and at most 7 of the FSHD-induced genes. However, all “FSHD-Hi” nuclei express at least 6 of these genes with most coexpressing at 10 and up to 18 genes (Figure 3C). To determine expression profiles of *DUX4*-detected nuclei, we examined genes coexpressed with *DUX4*. We found that *DUX4* was coexpressed with 20 FSHD-induced genes including two transcription factors, *LEUTX* and *ZSCAN4*, which have been reported as DUX4 targets in FSHD (Figure S7) [20, 22]. *DUX4* and *ZSCAN4* were expressed in all three *DUX4*-detected nuclei while *DUX4* and *LEUTX* were only simultaneously expressed in one *DUX4*-detected nuclei. FSHD-induced genes coexpressed in all three *DUX4*-detected nuclei include *KHDC1L*, *SLC34A2*, *PRAMEF9*, *RFPL4B* and *ZSCAN4*, while genes like *TRIM49*, *MBD3L2*, *MBD3L3* and *MBD3L5* are coexpressed with *DUX4* in two of the *DUX4*-detected nuclei. Additionally, the nucleus with *DUX4*, *LEUTX* and *ZSCAN4* also expressed *KDM4E*, *TRIM43*, *TRIM43B*, *MBD3L3*, *MBD3L5*, and *RFPL2*. Taken together, the genes expressed in the *DUX4*-detected nuclei may represent early targets of DUX4 which initiate a pathogenic gene regulatory network (Figure S7). Indeed, 18 of the 20 FSHD-induced genes expressed in at least one *DUX4*-detected nucleus are bound by DUX4 or in a gene cluster which is bound by DUX4 (i.e. MBD3Ls) [20, 22].

Interestingly, we observed the expression of DUX4 paralogs expressed in FSHD2 myotube nuclei. *DUXA* was expressed exclusively in the “FSHD-Hi” nuclei population while *DUXB* was only expressed in “FSHD-Lo” nuclei. In addition, *CHI3L1* was significantly higher in “FSHD-Lo” nuclei compared with “FSHD-Hi” myotube nuclei, while *CYTL1* was expressed exclusively in “FSHD-Lo” myotube nuclei (Figure 4B, 4C, S8) and in FSHD2 myoblasts (Figure 2A, 4C). To identify correlated gene regulation between the populations, we used the distinct expression profiles of *DUX4*, *DUXA*, *DUXB*, *CHI3L1* and *CYTL1* to mark the three FSHD2 myotube nuclei populations mentioned above and to observe their coexpression patterns with FSHD-induced genes (Figure 3D). We found that 19 FSHD-induced genes were coexpressed in both “FSHD-Hi” and “FSHD-Lo” populations, including reported DUX4 targets *LEUTX*, *ZSCAN4*, *MBD3L2*, *TRIM43*, *KHDC1L* and *CCNA1* [4, 20, 24], indicating that they may perform as a core set of responsive and interactive genes during FSHD progression (Figure 3D). We observed that 15 FSHD-induced genes were coexpressed only in the “FSHD-Hi” population, such as *KDM4E*, *PRAMEF1*, *PRAMEF11*, *PRAMEF12* and *SLC34A2*, and 9 genes were exclusively coexpressed in the “FSHD-Lo” population. As expected, *CYTL1* and *CHI3L1* were more frequently coexpressed in “FSHD-Lo” compared with other nuclei (Figure 3D). “FSHD-Hi” and “FSHD-Lo” have distinct coexpression patterns that indicate different cell states. Within the “FSHD-Hi” nuclei a large number of the FSHD-induced genes are coexpressed with transcription factors, such as *LEUTX* and *DUXA*, but not *DUX4*. Taken together, two identified patient myotube nuclei populations, “FSHD-Hi” with a small set of *DUX4*-detected nuclei and “FSHD-Lo”, exhibit distinct co-expression patterns of FSHD-induced genes including DUX4-target TF genes.

**Figure 4.**
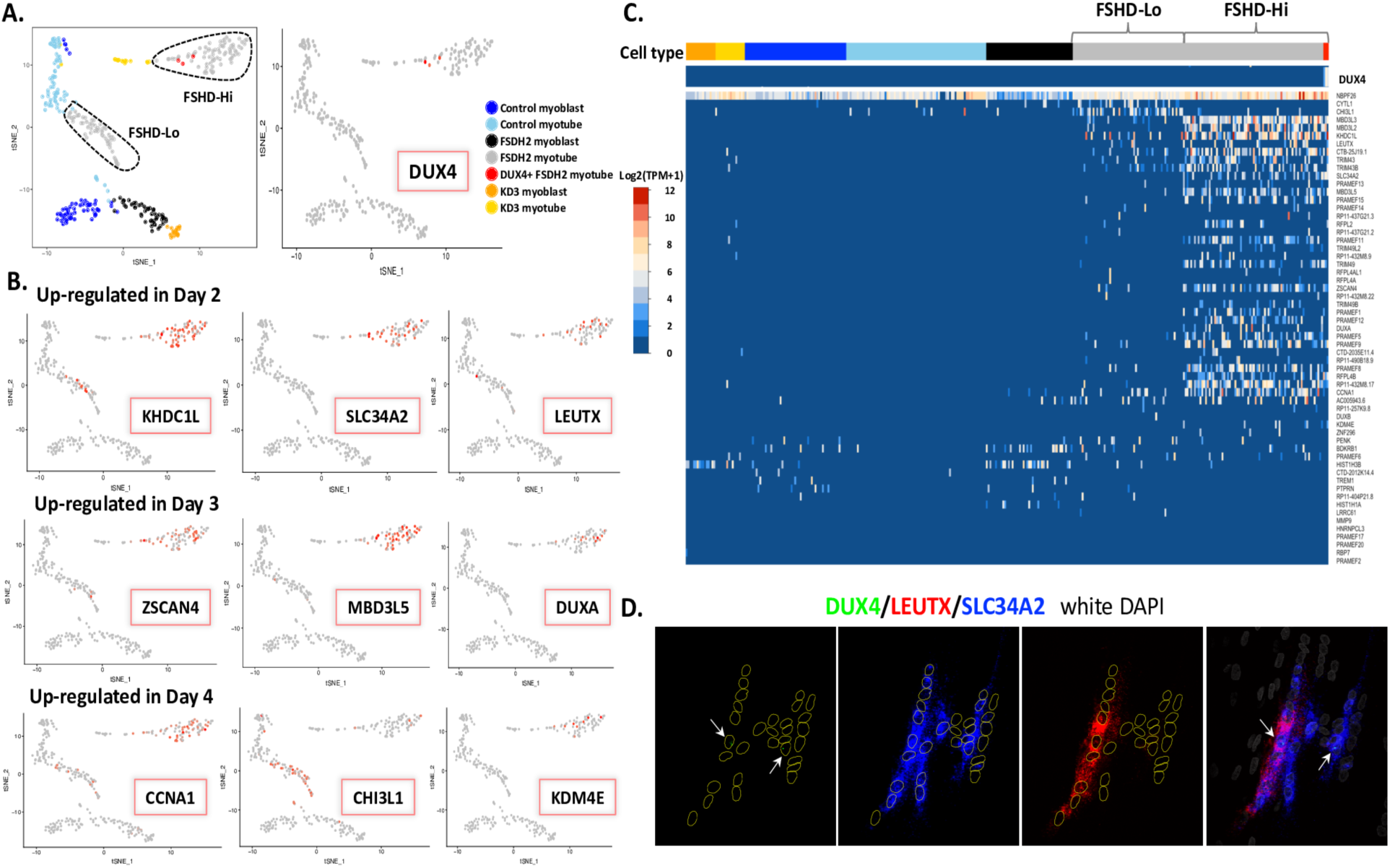
FSHD2 myotube nuclei show heterogeneous expression of *DUX4* and DUX4-associated FSHD-induced genes. **(A)** t-Distributed Stochastic Neighbor Embedding (t-SNE) plot of 349 single cells/nuclei RNA-seq data. Cell types are labeled by color, and the three *DUX4*-detected FSHD2 myotube nuclei are labeled in red. **(B)** Cells/nuclei are colored by the expression level of up-regulated genes in FSHD2 identified in time-course analysis. Genes are selected based on the day starting activated. Cells/nuclei with expression of specific genes are colored in red. **(C)** Gene expression heatmap of 59 FSHD-induced genes in FSHD2 that are up-regulated at day 3 or later stages. Cells and nuclei are ordered by cell types and labeled by distinct color in the annotation bar. High expression is shown in red and low expression is shown in blue. **(D)** RNA FISH of DUX4, LEUTX and SLC34A2 in FSHD2 myotubes at day 3 of differentiation. DUX4, green; LEUTX, red; SLC34A2, blue; DAPI, white.

### Heterogeneity of FSHD-induced gene expression in FSHD2 myotube nuclei

In order to further characterize the heterogeneous expression of *DUX4* and DUX4-associated FSHD-induced genes, we performed tSNE analysis on single-cell and single-nucleus RNA-seq samples. Similar to the PCA in Figure 3B, “FSHD-Hi” nuclei clearly separated from “FSHD-Lo” and control myotube nuclei (Figure 4A). Furthermore, we found that *DUX4* and FSHD-induced genes were often not expressed in the same nuclei (Figure 4A, 4B, S8-S11). To determine whether FSHD-induced gene expression was transient at the single nucleus level or persisted as expected from the pooled time-course (Figure 1C), we examined the expression of sets of genes significantly upregulated starting at days 2, 3 and 4 in our single-cell/single-nucleus data (Figure 4B). We found that genes significantly upregulated starting on days 2 and 3 in our pooled time course analysis were expressed in a higher proportion of nuclei and with higher expression levels in “FSHD-Hi” nuclei compared with other nuclei. We also observed this when looking at the expression of all 59 FSHD-induced genes in single cells and nuclei (Figure 4C). Interestingly, most of the genes with significant upregulation starting on days 0, 4 or 5 of differentiation were sparsely expressed in “FSHD-Hi” nuclei, and some had higher enrichment in “FSHD-Lo” nuclei (Figure 4C). For example, *CHI3L1*, significantly upregulated starting on day 4, was highly enriched in “FSHD-Lo” nuclei. We also observed that *CYTL1*, which was significantly more highly expressed in FSHD starting at day 0, was exclusively expressed in “FSHD-Lo” nuclei. To sum up, in “FSHD-Hi” we see that the genes which are upregulated beginning at day 2 are expressed along with those from day 3 in our day 3 myotubes, implying persistence of upregulation as seen in the pooled time-course. However, the “FSHD-Lo” myotubes show random expression of genes found to be upregulated very early or late in the pooled time-course, indicating the disease-related responses are more heterogeneous in “FSHD-Lo” than “FSHD-Hi” nuclei.

To substantiate the relationship between *DUX4*-detected nuclei and other nuclei from the same or different myotubes, we performed RNA FISH on *DUX4* and two representative FSHD-induced genes, *LEUTX* and *SLC34A2*, in 3-day differentiated FSHD2-2 myotubes. Probes were designed to hybridize to the two regions unique to the *DUX4fl* transcript to ensure the specificity. We observed that ∼7% of myotubes have at least 1 *DUX4*-detected nucleus, and that *DUX4*-positive myotubes contain on average 2 *DUX4*-detected nuclei (among on average 15 nuclei per myotube). While the *DUX4* transcript is often detectable only in the nucleus, *LEUTX* and *SLC34A2* mRNA transcripts are abundantly present in the cytoplasm as well as in multiple nuclei suggesting that the DUX4 protein activates their transcription in multiple nuclei of a single myotube (Figure 4D). Furthermore, the *DUX4* transcript signal is very low or undetectable in some myotubes while *LEUTX* and *SLC34A2* transcripts are detected (Figure 4D, S12). These results strongly suggest that FSHD-induced genes can persist even after *DUX4* transcription has ceased, which explains the long-standing observation in previous studies that FSHD-induced gene expression is easier to detect in patient muscle cells than the *DUX4* transcript itself.

### “FSHD-Hi” nuclei form a distinct branch on the pseudotime of single-cell and single-nucleus differentiation

We performed a pseudotime analysis using Monocle [27] in order to understand whether the cells expressing FSHD-induced genes followed a distinct developmental trajectory (Methods). We reordered 349 single-cells/single-nuclei based on pseudotime (Figure S5A), which recapitulated the differentiation trajectory from myoblast to myotube and identified additional subpopulations in myotube nuclei compared with myoblast cells (Figure 5A). Interestingly, the “FSHD-Hi” nuclei population with three *DUX4*-detected myotube nuclei formed a homogenous cluster (branch 7, with only one KD3 myotube nucleus) at one end of the pseudotime and showed upregulation of the FSHD-induced genes that we had identified in our pooled RNA-seq differentiation time-course (Figure 5A). Nuclei from the “FSHD-Lo” population mixed with many control myotube nuclei (branches 2 and 3), while some located on the same branch as myoblast cells (branch 1) in terms of pseudotemporal position (Figure 5A) indicating that the heterogeneity in this population might be in part caused by a less advanced differentiation status. We further characterized the expression of genes induced late in FSHD2 differentiation (cluster 4, Figure 2D) in single cells/nuclei (Figure 5C). Late-induced genes were more highly enriched in FSHD2 myotube nuclei compared with others and were almost entirely not expressed in myoblast cells (Figure S13A). Genes induced at the beginning of differentiation (cluster 1, Figure 2B), however, were expressed in both myoblast cells and myotube nuclei (Figure 5B, S13B). These results further confirm the importance of later-stage induced genes in distinugishing between control and disease phenotypes. As mentioned above, genes induced late in FSHD2 differentiation (cluster 4, Figure 2D) were detected in the FSHD2 single myotube nuclei include known DUX4 targets such as *ZSCAN4*, *MBD3L2*, *KHDC1L*, *LEUTX* [4, 20, 24], as well as DUX4 paralogs *DUXA* and *DUXB*. We found these DUX4 targets, including *DUXA* but not *DUXB,* were also coexpressed in the “FSHD-Hi” nuclei without *DUX4* detected (branch 7) (Figure 3D, S14) strongly suggesting that these nuclei are responding to the presense of the DUX4 protein in the same myotubes even though *DUX4* is not being transcribed in the same nuclei.

**Figure 5.**
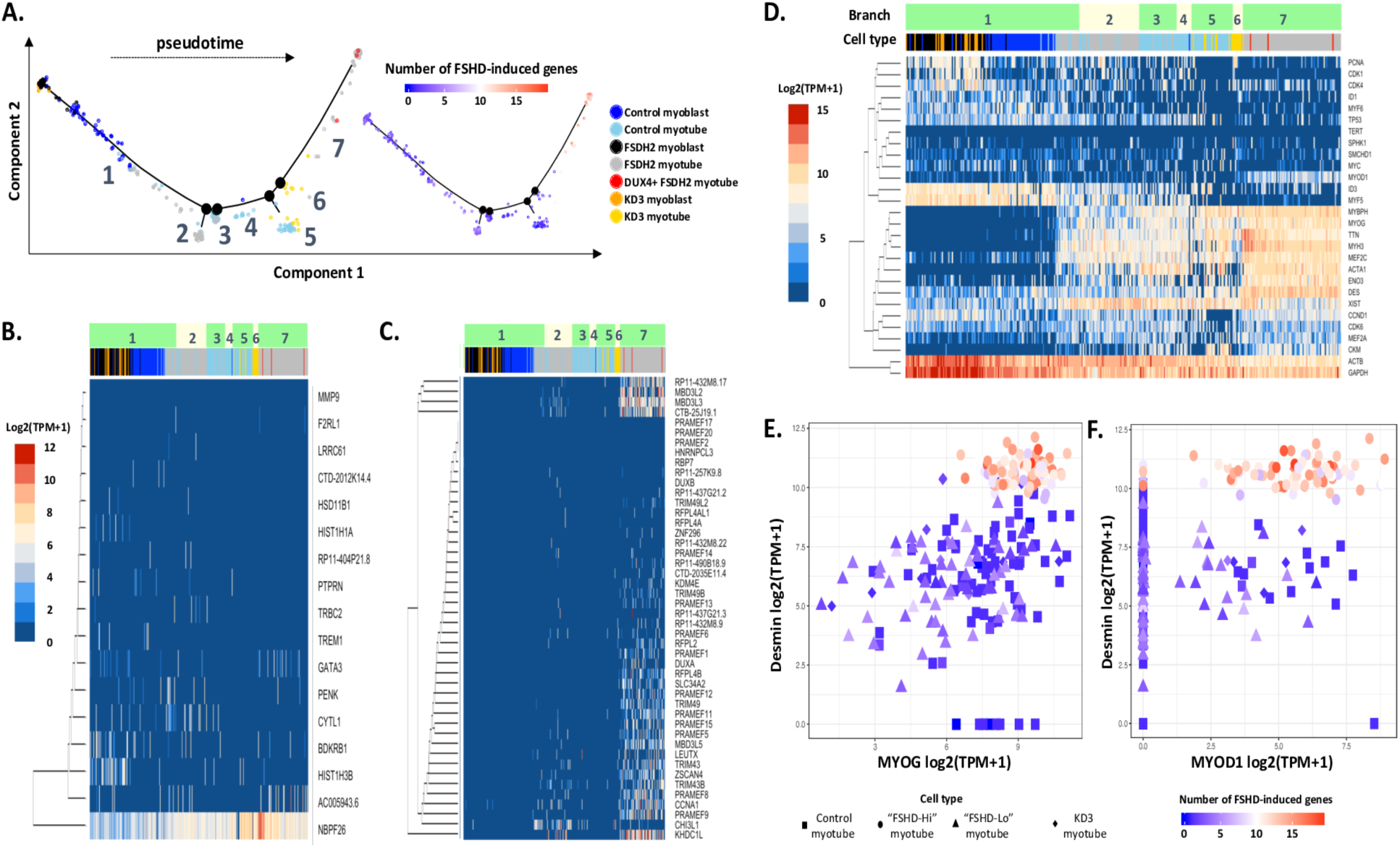
Pseudotime ordering separates “FSHD-Hi” nuclei from all other cells and nuclei. **(A)** Pseudotemporal ordering of 349 single cells/nuclei using independent component analysis by Monocle. Seven branches (1-7) are identified to separate cells/nuclei into subpopulations. Cell types are labeled by color. Nuclei on branch 7 are highly enriched in DUX4 targets. **(B)** Gene expression heatmap of the 17 genes from cluster 1 (Figure 2B) in single cells/nuclei following pseudotime of differentiation. Cells/nuclei are ordered by pseudotime and cell types are labeled in the annotation bar. **(C)** Gene expression heatmap of the 46 genes from cluster 4 (Figure 2D) in single cells/nuclei following pseudotime of differentiation. Cells/nuclei are ordered by pseudotime and cell types are labeled in the annotation bar. **(D)** Gene expression heatmap of myogenic markers in single cells/nuclei (ordered by pseudotime generated by Monocle). **(E)** Scatterplot of gene expression between *MYOG* and *desmin* in myotube nuclei. **(F)** Scatterplot of gene expression between *MYOD1* and *desmin* in myotube nuclei. Cell types are labeled by shape. Single nuclei are colored by the number of FSHD targets in FSHD2 during differentiation.

Although DUX4 and FSHD-induced genes are thought to inhibit myogenesis in FSHD2 [28], the pseudotemporal positions of cells and nuclei suggest that myotube maturation may also affect the expression of *DUX4* and its targets. We found that markers of myotubes, such as *TTN* and *MYBPH*, have higher expression in branches 5, 6 and 7 compared with the ones in branches 2 and 3 (Figure 5D, S15A) with highest expression in branch 7. However, *ACTA1* was highly expressed in branches 2, 3 and 7 but not in 5 and 6, while *CKM* was highly expressed in branches 5 and 7 but not others (Figure 5D, S15A). Interestingly, we found that *desmin* was also significantly more highly expressed in branch 7 (“FSHD-Hi” nuclei) than other branches (Figure 5D, S15B). By comparing between branches, we found that *desmin*, *MYOG* and *ACTA1* were significanly upregulated in branch 7 (“FSHD-Hi”) (Figure 6A) while they showed similar expression levels between branches 2 and 3 (“FSHD-Lo” with some control nuclei) and branches 5 and 6 (control and KD3 nuclei). We also observed that “FSHD-Hi” myotube nuclei expressing more FSHD-induced genes had higher expression of both *desmin, MYOG* (Figure 5E) and *ACTA1* (Figure S15C) but had similar expression levels of *MYOD1* (Figure 5F) and *CKM* (Figure S15D) compared with other myotube nuclei. Given that the “FSHD-Hi” nuclei in branch 7 expressed more *desmin* than the “FSHD-Lo” nuclei in branches 2 and 3, we propose that the maturation state of the myotube is closely linked to *DUX4* and FSHD-induced genes expression and that high *desmin* expression appears to signify the “FSHD-Hi” state of patient myocytes.

**Figure 6.**
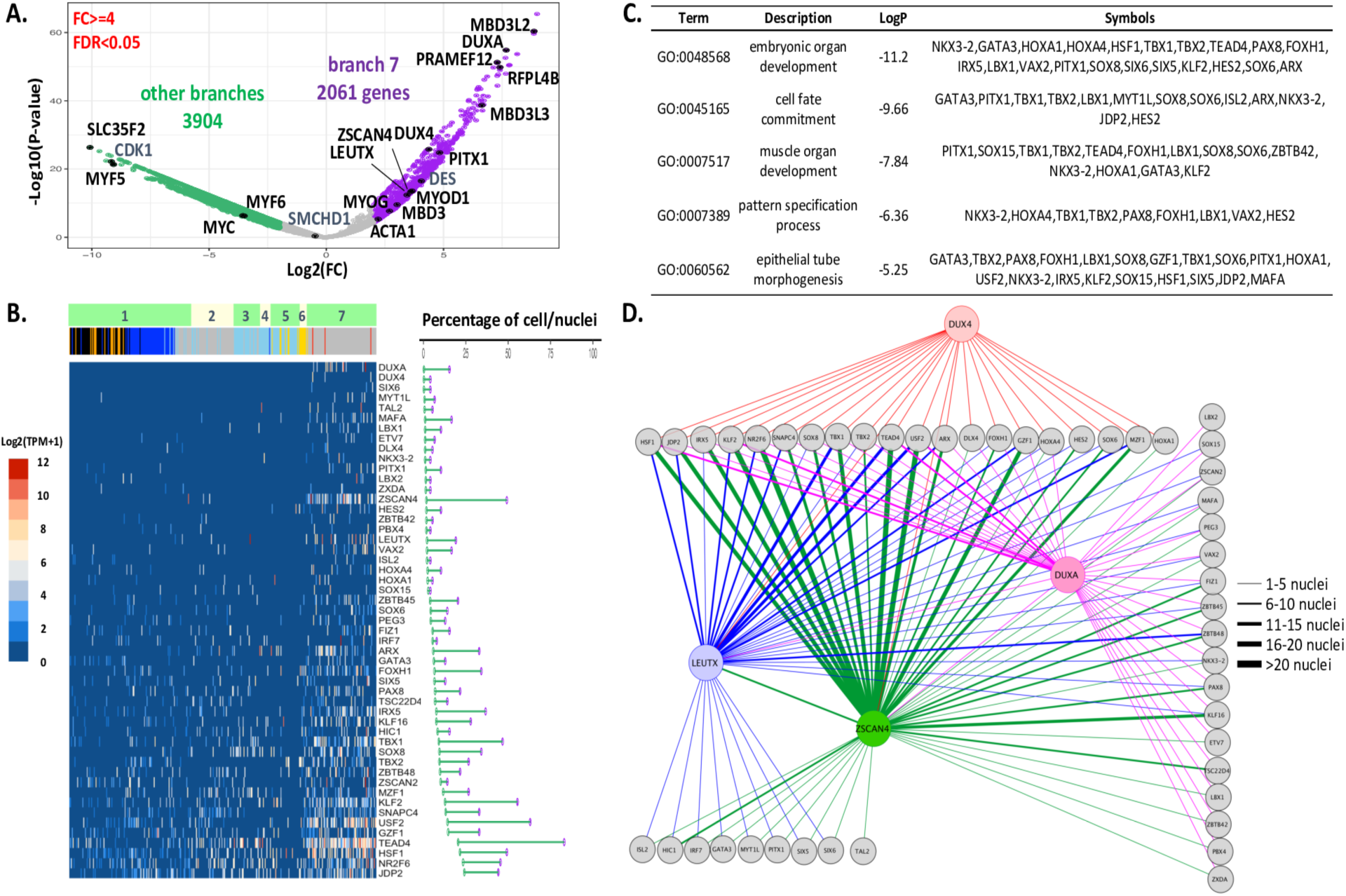
Identification of TFs significantly enriched in “FSHD-Hi” myotube nuclei. **(A)** Differential expression analysis between branch 7 and all other branches from Figure 5A. Differentially expressed genes are colored for each branch (FDR < 0.05, log2 (fold change)>=2). **(B)** Gene expression heatmap of 51 up-regulated TFs in branch 7. TFs are sorted by the number of nuclei expressing these genes in other branches. Comparison of the number of nuclei expressing each TFs in both branch 7 (purple dot) and others (green dot) is shown on the right of the heatmap. **(C)** Significant gene ontology terms associated with 51 up-regulated TFs in branch 7 (FDR < 0.05). **(D)** Coexpression network between DUX4, DUXA, LEUTX, ZSCAN4 and other TFs in branch 7.

### Identification of TFs as potential regulators downstream of DUX4 in FSHD2

**“**FSHD-Hi” myotube nuclei are highly enriched in FSHD-induced genes compared to “FSHD-Lo” or control myotube nuclei. Interestingly, more than half of the FSHD-induced genes that are coexpressed in “FSHD-Hi” are also coexpressed in the “FSHD-Lo” population, providing evidence that even at a low level “FSHD-Lo” cells are undergoing disease-specific changes (Figure 3D). Of the FSHD-induced genes that are coexpressed in *DUX4*-detected nuclei, 95% are also coexpressed in other “FSHD-Hi” nuclei, supporting the notion that DUX4 protein diffuses to and is exerting its regulatory effects in multiple nuclei within the same myotube (Figure S7, S14). Interestingly, however, we found that a unique subset of FSHD-induced targets are coexpressed with either *DUXA*, *LEUTX*, and/or *ZSCAN4* only in “FSHD-Hi” nuclei without *DUX4* detected, indicating the varied transcriptional responses in different nuclei (Figure S14). Downstream TFs of DUX4 may interact with these FSHD-induced genes and play important roles in mediating the pathogenic program.

To further elucidate the possible regulators of this program, we performed differential expression analysis between nuclei in branch 7 (“FSHD-Hi”) and other branches. We identified 2,061 genes significantly upregulated in branch 7 including myogenic differentiation markers and multiple FSHD-induced genes (Figure 6A, 6B). *MYOD1*, *MYOG* and *ACTA1* were more highly expressed in branch 7 “FSHD-Hi” nuclei while a cell cycle gene, *CDK1*, was downregulated (Figure 5D, 6A). The myoblast marker *MYF5* was one of the top upregulated TFs in the other branches compared to branch 7 (Figure 6A). Additionally 50 TFs were upregulated in branch 7 which did not include *DUX4* but did include previously reported TFs activated in FSHD, such as *LEUTX*, *ZSCAN4*, *PITX1*, and *DUX4* [4, 20, 24, 29]. We observed several TFs, such as *DUXA*, *GATA3*, *TEAD4*, *LBX1*, *TBX1*, *KLF2*, *HOXA1*, *HOXA4*, *FOXH1* and *NKX3-2*, associated with embryonic organ development and muscle organ development upregulated in most of the nuclei in branch 7 but in less than 25% of cells or nuclei in other branches (Figure 6B, 6C). These results indicate the activation of a distinct gene regulatory network in branch 7 “FSHD-Hi” nuclei.

To gain insight into a possible network of gene regulation downstream of DUX4 activation, we analyzed the coexpression patterns of these TFs in “FSHD-Hi” myotube nuclei and found similar coexpression patterns for the TFs as was found for the FSHD-induced genes (Figure S14). The majority of TFs coexpressed with *DUX4* in *DUX4*-detected nuclei were also coexpressed with *DUXA* in other “FSHD-Hi” nuclei, and many TFs were coexpressed only in these nuclei (Figure 6D, S14). Twenty TFs were coexpressed with *LEUTX* and *ZSCAN4* in both *DUX4*-detected and other “FSHD-Hi” nuclei, and these TFs were significantly enriched in embryonic organ development (p = 1 × 10^−8.19^) and chordate embryonic development (p = 1 × 10^−5.89^), including *HOXA4*, *HSF1*, *TBX1*, *TBX2*, *TEAD4*, *FOXH1*, *IRX5*, *SOX8*, *KLF2*, *ARX*, *HES2*, *HOXA1*, *SOX6* and *MZF1*. HES2 is involved in the PI3K-Akt signaling pathway and is known to regulate immue cell development as well as repressing myogenesis by Notch signaling [30, 31]. SOX6 and MZF1 have been reported to regulate the FSHD region genes *FRG1* and *FRG2KP* [32]. Another 27 TFs were only expressed in “FSHD-Hi” nuclei with no *DUX4* detected and were enriched in neuron fate specification (p = 1 × 10^−5.18^) and muscle organ development (p = 1 × 10^-3.01^). Of these, ZBTB48 is known to directly bind to telomeres and to regulate their maintainance by limiting their elongation [33]. ZSCAN2, a testis-expressed gene, is also upregulated in “FSHD-Hi” nuclei, which is consistent with previous studies on DUX4-activated genes involved in germ cell and early development [4, 34]. We found that TFs which regulate muscle atrophy wasting, such as PEG3 [35] and ZBTB42 [36], were also upregulated in “FSHD-Hi” nuclei. In addition, *PITX1* and 7 others were coexpressed with *LEUTX* and *ZSCAN4* but not *DUXA* or *DUX4* in “FSHD-Hi” myotube nuclei. Taken together, single nucleus analysis revealed that athough a set of TFs are upregulated in “FSHD-Hi” nuclei as a whole, *DUX4*-detected nuclei and other “FSHD-Hi” nuclei have distinct sets of TFs coexpressed raising the possibility that the nuclear environment which permits or does not permit *DUX4* expression may differentially affect the expression of other genes, invoking activation of divergent gene regulatory networks.

A similar single-cell RNA-seq study was published recently that also identified a small population of *DUX4* transcript-positive cells in both FSHD1 and FSHD2 patient-derived primary myocytes [37]. In their study, however, myoblast differentiation was induced but myotube formation was artificially blocked by the use of a calcium chelator [37]. This is in contrast to our study, in which we examined nuclei from unperturbed myotubes using snRNA-seq. Importantly, our approach enables us to = address how *DUX4* expression, even in a single nucleus, results in target gene activation in other nuclei in the same myotube (due to the DUX4 protein spreading) under native condition to distinguish the “FSHD-Hi” and “FSHD-Lo” population of cells. We analyzed the expression of 67 DUX4 target genes used in Heuvel, et al. [20, 37] in our “FSHD-Hi” and “FSHD-Lo” myotube single nucleus populations. All “FSHD-Hi” nuclei and about 3.3% of “FSHD-Lo” nuclei highly expressed at least 5 of these genes, which is much higher than that in single cell myocyte data (0.2-0.9%) [37] (Figure S16). Of note, we identified that 2.12% of nuclei in our primary FSHD2 myotubes expressed high levels of *DUX4*, which is higher than the percentage reported in single cell myocytes (0.2-0.9%) [37] (Figure S16A). It is currently unclear whether blocking myotube fusion interferes with the normal course of myotube differentiation and affects frequency of *DUX4* expression. Thus, our snRNA-seq analysis captured the extent of target gene expression by the limited expression of *DUX4* in patient myotubes. Our higher-sensitivity allowed us to identify and distinguish the behaviors of the “FSHD-Hi” and “FSHD-Lo” populations, possibly representing two different states of patient myotubes.

## Discussion

Using our pooled RNA-seq time-course, we discovered two sets of FSHD-specific genes that exhibit distinct expression dynamics: 17 genes that are differentially upregulated as early as the myoblast stage (“early-induced”) and a second set of 46 genes that are significantly upregulated starting at day 2 and stably activated by day 3 during FSHD2 myoblast differentiation (“late-induced”). We defined 59 genes that are stably activated by day 3 of differentiation as “FSHD-induced genes” (defined in results section). Previous studies have confirmed many genes as downstream targets of DUX4 by transcriptome and ChIP-seq analysis [20, 24]. We found that these known DUX4 targets show higher enrichment in late-induced FSHD associated genes in our study, and that other FSHD-induced genes from the same late-induced cluster may be novel DUX4 targets. Our single-cell and single-nucleus RNA-seq analysis on control and FSHD2 myoblasts and 3-day differentiated myotubes, respectively, confirmed that FSHD2 myotube nuclei have significantly higher enrichment of FSHD-induced genes than myoblasts and control myotube nuclei. Importantly, we were able to identify *DUX4* transcript-positive nuclei, which was not detected in our pooled RNA-seq. We further identified two populations of FSHD2 myotube nuclei, “FSHD-Hi” and “FSHD-Lo”, based on the expression levels of FSHD-induced genes and showed that *DUX4*-detected nuclei belong to “FSHD-Hi”. Late-induced genes showed higher expression levels in “FSHD-Hi” than in “FSHD-Lo”, and a higher proportion of “FSHD-Hi” nuclei express these genes. Although both “FSHD-Hi” and “FSHD-Lo” were collected on the same day, “FSHD-Lo” shows relatively earlier differentiation status based on the time-course data and pseudotime analysis. Interestingly, however, both early-induced genes and most of the genes that are induced later on days 4 and 5 were sparsely expressed in “FSHD-Hi” nuclei, and some of were more highly enriched in “FSHD-Lo” nuclei. Therefore, our results reveal that the expression of *DUX4* and its downstream targets is closely linked to the precise differentiation status of myotube and that less mature myotubes have more heterogenous responses of FSHD-induced genes.

While we only detected a small set of *DUX4* transcript-positive nuclei, the proportion (3/79, 3.8%) is higher than the reported 0.05% (1/200 in myotube nuclei) [8, 10, 16], indicating that expression levels of *DUX4* could vary between individuals or be very sensitive to growth and differentiation conditions. Furthermore, we found that the expression of *DUX4* in a single nucleus was higher than stated in previous population studies [8, 38] as we detected *DUX4* expressed at 11.24 TPM, 34.15 TPM and 68.49 TPM in individual nuclei. Considering that DUX4 target genes were most stably upregulated at day 3 of differentiation in our time-course, our results strongly suggest that maturation status has a substantial impact on the expression of *DUX4* and its target genes. Studies have previously found that *DUX4* expression caused defects in muscle development and even led to cell death [14,15]. One possible hypothesis integrating these observations is that *DUX4* only activates the pathogenic FSHD gene network when expressed during a critical time window in myogenesis. Outside of this window transient *DUX4* expression cannot activate these same targets possibly due to lack of necessary cofactors or epigenetic states. Pathogenesis then would have to be regulated through a downstream network of gene dysregulation. *DUX4* is known to be temporally activated as a key regulator during human early embryonic development during which it activates genes involved in zygotic genome activation (ZGA) [11]. Indeed, dysregulation of *DUX4* in muscle has been shown to activate germline-specific genes [4, 22, 34]. The widespread, self-supported network of gene regulation necessary for ZGA may be what proves toxic for terminally differentiated muscle. Further experiments are required to validate this proposed mechanism.

Small populations of DUX4-positive myotubes are thought to drive pathogenesis in FSHD [2-4], which may explain the slow progress of the disorder. It is still unclear whether DUX4-negative patient muscle cells are normal or contribute to disease pathogenesis. Our high-resolution single-cell and single-nucleus dataset is the first to observe the endogenous expression of *DUX4* in a small number of FSHD2 myotube nuclei. We found that these nuclei cluster with a much larger set of FSHD2 nuclei expressing multiple FSHD-induced genes which we define as “FSHD-Hi”. *In situ* RNA FISH analysis showed that *DUX4* target genes are expressed in the same myotube together with a few *DUX4*-detected nuclei. Thus, our results clearly demonstrate that *DUX4* expressed even in one nucleus is sufficient to activate DUX4 target gene expression in multiple nuclei in a single FSHD myotube. This is consistent with previous studies showing that both overexpressed recombinant and endogenous DUX4 protein diffuses to multiple nuclei within a myotube [16, 22]. We also observe a distinct population of FSHD2 nuclei (termed “FSHD-Lo”) which express fewer FSHD-induced genes than “FSHD-Hi” but more than control myotube nuclei. “FSHD-Hi” and “FSHD-Lo” nuclei separated into distinct populations on the pseudotime, and both of populations clearly separated from control myotube nuclei. “FSHD-Lo” nuclei thereby represent distinct myotube population from “FSHD-Hi” nuclei. Importantly, our results indicate that even these *DUX4*-negative patient muscle cells exhibit disease-specific changes and suggest that they may also contribute to the disease.

While we detected FSHD-induced genes in both “FSHD-Hi” and “FSHD-Lo” myotube nuclei, many of them are coexpressed in only a subset of cells, demonstrating heterogeneous expression patterns at the single-nucleus level. Interestingly, we find distinct coexpression patterns involving *DUXA*, *LEUTX*, *ZSCAN4* and other FSHD-induced genes, such as 7 *PRAMEF* genes and *TRIM49L2* in “FSHD-Hi” myotube nuclei. Although DUXA is a DUX4 paralog [25] and has been identified as a DUX4 target gene in human patient muscle cells [20], almost no study reports its specific functions in FSHD2 development. Similar to DUX4, DUXA is a transcription factor with two homeobox domains, and it binds to a 10bp motif similar to DUX4 [39]. Given that *DUX4* and *DUXA* were not coexpressed in FSHD2 myotube nuclei and that *DUXA* was frequently coexpressed with FSHD-induced genes, we propose that, similar to DUX4, DUXA may drive an FSHD-specific pathogenic program by binding and activating a pool of downstream targets, therefore reinforcing DUX4-induced gene network in patient myotube nuclei.

Interestingly, we found novel gene markers, *CYTL1* and *CHI3L1*, which distinguish “FSHD-Lo” myotube nuclei from “FSHD-Hi” nuclei. Although *CHI3L1 (Chitinase-3-like-1)* was expressed in a few “FSHD-Hi” nuclei, the proportion and expression levels were significantly higher in the “FSHD-Lo” population. *CYTL1 (Cytokine-like 1)* was found only in “FSHD-Lo” nuclei. *CHI3L1* is located just upstream of *MYOG* and is more highly expressed in myotubes than myoblasts [40]. It has been shown to be expressed by exercised mytoubes and to stimulate myoblast differentiation [41]. Additionally, CHI3L1 is known to interfere with TNFα-mediated inflammation in muscle cells [42] but is implicated in pro-inflammatory response [43] and in fast developing DMD, in which inflammation exacerbates muscle dystrohpy [44]. CYTL1 has been shown to significantly prevent inflammatory arthritis [45] and has been proposed to have anti-inflammatory effects [46]. CHI3L1 and CYTL1 have also been found to be upregulated in FSHD muscle biopsies with low expression of TRIM43, LEUTX, PRAMEF2, KHDC1L, similar to our “FSHD-Lo”. These tissues have mild pathology and normal MRIs with no signs of active disease, but show evidence of activation of the complement system [47]. Given the frequent coexpression of *CHI3L1* and *CYTL1* in “FSHD-Lo” nuclei, an antagonistic interaction between these two genes may possibly help to maintain homeostasis and further alleviate the pathological effects of FSHD-induced genes in the myotube.

Although we have detected a small set of FSHD2 myotube nuclei that express high expression of *DUX4*, DUX4 target genes are expressed in a larger fraction of myotube nuclei. The target gene products are also expected to diffuse through the cytoplasm of the myotubes and to alter the downstream gene expression profiles in multiple nuclei to further contribute to FSHD2-specific pathogenesis. We found a set of TFs that are specifically upregulated in “FSHD-Hi” myotube nuclei and that more than half of these are frequently coexpressed with *LEUTX* and *ZSCAN4*, the two known DUX4 target TFs in “FSHD-Hi” nuclei. These “FSHD-Hi”-upregulated TFs are associated with germ cell and early development, and some of them have been reported to promote muscle atrophy wasting [35, 36]. These TFs may be novel DUX4 target genes or further downstream DUX4-responsive genes which regulate a network of genes in FSHD-Hi nuclei eventually leading to the myotoxicity and dystrophy by amplifying or reinforcing the effects of the sporadic and transient expression of *DUX4* through self-sustained gene dysregulation. Although we did not find significant upregulation of *P53* in our “FSHD-Hi” population, the P53 upstream activator, HIC1 [48, 49], and the P53 signaling mediator, PEG3 [35], were upregulated in “FSHD-Hi” nuclei, and *HIC1* was coexpressed with the DUX4 target *PITX1,* which is known to induce *P53* and further promote muscle atrophy by increasing inflammation and oxidative stress [50-53]. These observations challenge the assumption that endogenous *DUX4* expression is autonomously toxic to the cell and raises the possibility that DUX4 triggers coexpression of multitude of DUX4 target and responsive TFs that serve as mediators of FSHD. These TFs may coordinately regulate a network of genes resulting in sustainable activation of the cytotoxic program that leads to pathogenesis. If this is the case, therapeutics targeting DUX4 or *DUX4* expression may limit progression of FSHD to new tissue but may not stop muscle wasting in already disease-activated tissue.

Our time-course pooled RNA-seq and single cell/nucleus RNA-seq of primary control and FSHD2 myoblasts and myotubes revealed distinct dynamics of FSHD-specific gene expression during differentiation and demonstrated that all the patient cells exhibit disease-specific gene expression changes even in those with no detectable DUX4 as well as very low FSHD-induced genes expression. We provided direct evidence that DUX4 transcript expression even in a single nucleus can trigger a high expression of downstream target genes in the same myotube, but the expression pattern of downstream genes can vary between nuclei. Specific analysis of nuclei with detectable *DUX4* and/or FSHD-induced genes expression revealed co-upregulation of a specific set of transcription factors that may together contribute to the development of the FSHD phenotype.

## Methods

### Human myoblast culture and differentiation

Human control and FSHD2 myoblast cells from patient quadricep and tibia biopsies, and KD3 immortalized cells from abdominal muscles, were grown on dishes coated with collagen in high glucose DMEM (Gibco) supplemented with 20% FBS (Omega Scientific, Inc.), 1% Pen-Strep (Gibco), and 2% Ultrasor G (Crescent Chemical Co.) [21]. Upon reaching 80% confluence, differentiation was induced by using high glucose DMEM medium supplemented with 2% FBS and ITS supplement (insulin 0.1%, 0.000067% sodium selenite, 0.055% transferrin; Invitrogen). Fresh differentiation medium was changed every 24hrs.

### Pooled, single-nucleus and single-cell RNA-seq library preparation and sequencing

For pooled RNA-seq for the time-course, total RNA was extracted by using the RNeasy kit (QIAGEN). Between 19 and 38 ng of RNA were converted to cDNA using the SmartSeq 2 protocol [54]. Libraries were constructed with the Nextera DNASample Preparation Kit (Illumina). Single-cell and single-nucleus RNA-seq were performed according to [21] with the following modifications. Myotube single nuclei were isolated from mononucleated cells (MNCs) by washing a 6 cm dish once with trypsin, then adding trypsin for about 5 min until myotubes lifted off the plate and MNCs were still attached. Cells were initially pelleted at 2000 rpm for 2 min and resuspended in lysis buffer with 0.02% IGEPAL CA-630. Lysis was done for 3 minutes, filtered and spun at 4000 rpm for 1 minute. Nuclei were captured on medium IFCs (10-17 um) at a density between 340 and 640 nuclei/ul in a volume of 10 ul. Visual confirmation was aided with the LIVE/DEAD kit (Thermo Fisher Scientific), and cDNA was normalized to approximately 0.1 ng/ul for tagmentation and library prep. Libraries were quality-controlled prior to sequencing based on Agilent 2100 Bioanalyzer profiles and normalized using the KAPA Library Quantification Kit (Illumina). Libraries were sequenced on the Illumina NextSeq500 platform using paired-end 43bp mode with 15 million reads per sample for pooled libraries and paired-end 75bp mode with 1-3 million reads per sample for single-cell and single-nucleus libraries.

### RNA FISH (Fluorescent *in situ* hybridization targeting ribonucleic acid molecules)

FSHD2-19 myoblasts were seeded in micro-slide eight-well plates at ∼8×10^4^ cells per well, and differentiation was initiated ∼20hrs later. After 3 days of differentiation, cells were fixed with 10% neutral buffered formalin at room temperature for 30 min, and the RNA FISH experiments were performed using the RNAscope fluorescent Multiplex system (Advanced Cell Diagnostic Inc.) according to the manufacturer’s instructions. Probe-Hs-DUX4-C1, Probe-Hs-LEUTX-C2, and Probe-Hs-SLC34A2-C3 were used. Images were acquired using a Zeiss LSM 510 META confocal microscope.

### RNA-seq data processing

Raw reads from both pooled RNA-seq and single-cell and single-nucleus RNA-seq were mapped to hg38 by STAR (version 2.5.1b) [55] using defaults except with a maximum of 10 mismatches per pair, a ratio of mismatches to read length of 0.07, and a maximum of 10 multiple alignments. Quantitation was performed using RSEM (version 1.2.31) [56] by defaults with gene annotations from GENCODE v28, and results were output in transcripts per million (TPM). Myoblast cells were kept for downstream analysis if *desmin* expression was >=1 TPM, *MYOG* <1 TPM, number of expressed genes was more than 500 and expression level of *GAPDH* was higher than 100 TPM. Myotube nuclei were kept for downstream analysis if *MYOG* expression was >=1 TPM, number of expressed genes was more than 500 and expression level of *GAPDH* was higher than 100 TPM. We only kept cells and nuclei with a uniquely mapped efficiency higher than 45%. For differential gene expression analysis in differentiation time-course, protein coding and long non-coding RNA genes with at least 5 TPM in both replicates in at least one timepoint and with at least 1 TPM for both reps for both cell lines of the same disease and day were kept. Genes were TMM normalized using edgeR (version 3.18.1) [57] and log2-transformed for downstream analysis. Differential expression analysis per day during differentiation was done by using edgeR with FDR < 0.05. Clustering of differentially expressed genes across the time-course was done by using maSigPro [26]. For dimensionality reduction analysis in single-cell and single-nucleus RNA-seq data, incremental PCA analysis was performed by the *IncrementalPCA* function from scikit-learn [58], python 2. t-SNE analysis was performed by using Seurat [59]. Monocle (version 2.4.0) [27] was used to determine the pseudotime trajectory. Gene ontology analysis was done by using Metascape [60] with FDR<0.05. Gene coexpression networks were plotted by using Cytoscape [61].

### Data availability

All pooled-, single cell, and single nucleus RNA-seq data along with their associated meta data will be deposited in the GEO database.

## Acknowledgements

We thank the UCI GHTF for access to the Fluidigm C1. This work was supported in part by grants from the National Institutes of Health (AR071104 and AR071287 to K. Y. and A.M.).

## Supplementary file

**Figure S1.**
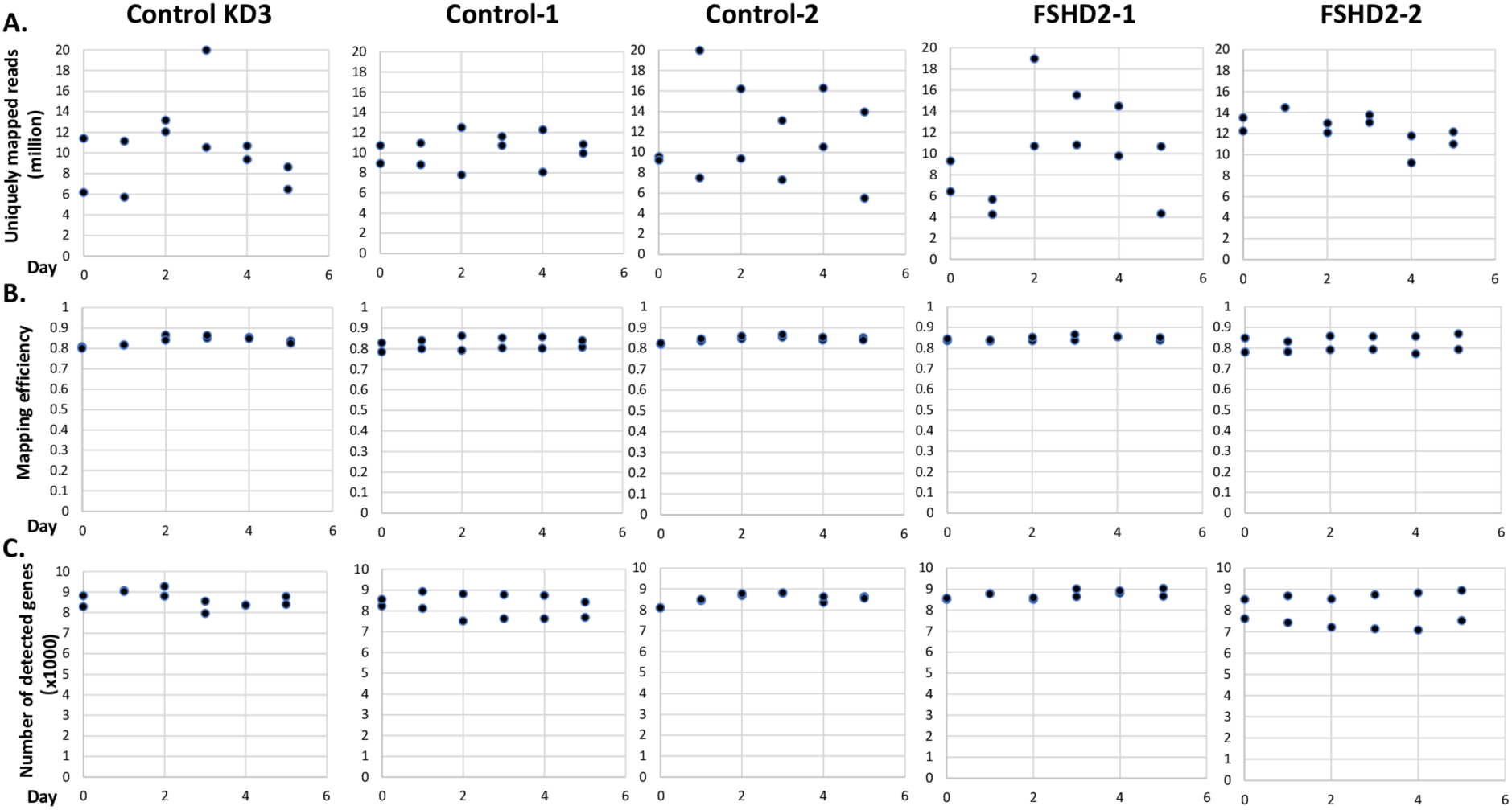
Quality metrics of RNA-seq time-course data. Control, FSHD2 and KD3 time-course quality metrics for **(A)** the number of uniquely mapped reads, **(B)** mapping efficiency, **(C)** the number of genes detected (TPM>=1).

**Figure S2.**
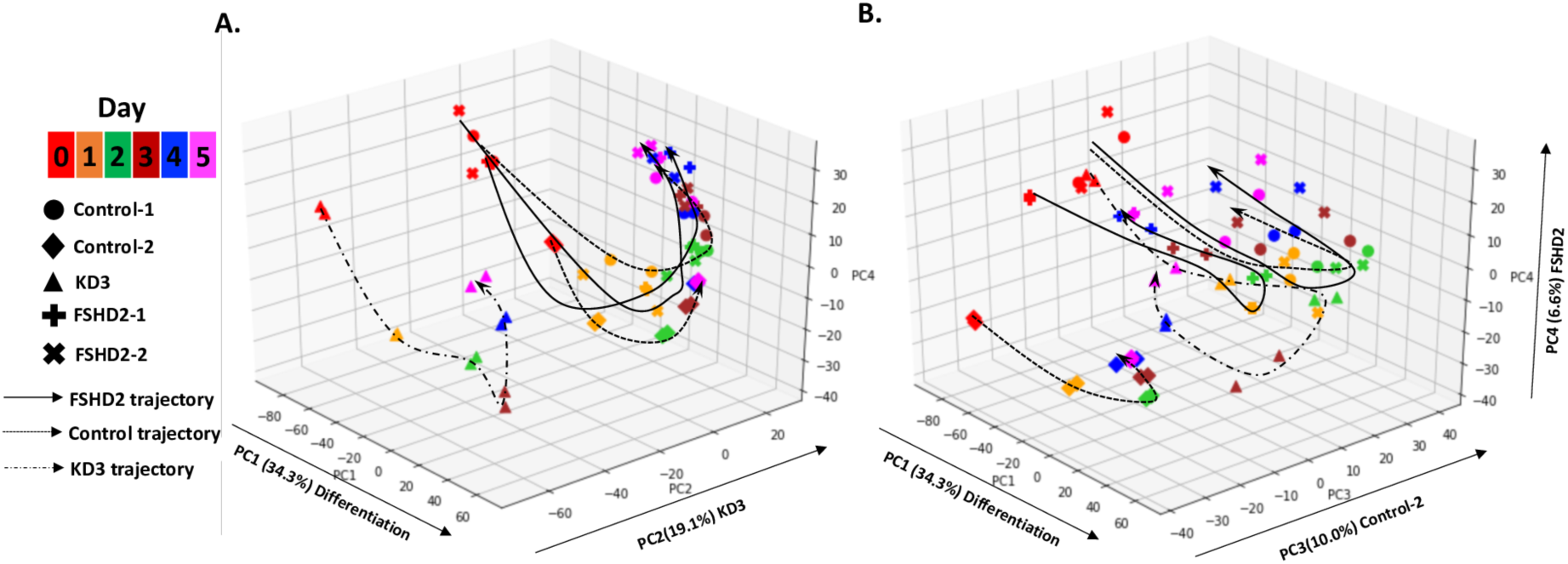
Principal component analysis (PCA) on control, FSHD2 and KD3 myoblast differentiation time-course. **(A)** PCA with PC1, PC2 and PC4. PC2 explains the expression variance between KD3 and others. **(B)** PCA with PC1, PC3, and PC4. PC3 explains the expression variance between Control-2 and all others. Gene expression level was measured each day for duplicates by using RNA-seq. Cell types are labeled by shape, and time-points are labeled by color.

**Figure S3.**
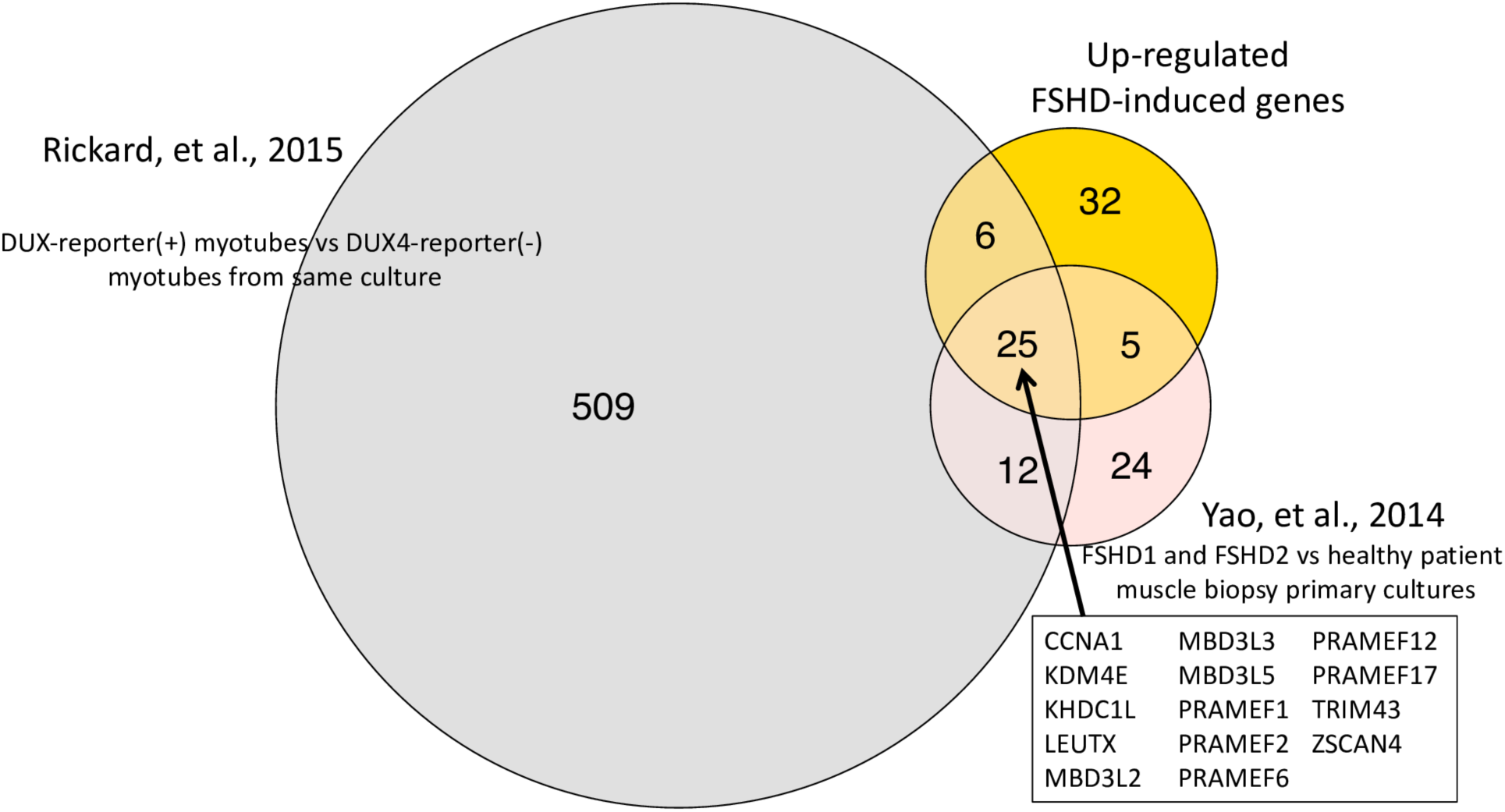
Venn diagram of FSHD-induced genes from this study and other published FSHD or DUX4 induced genes. Overlap of 68 genes upregulated during FSHD2 differentiation time-course from myoblasts to myotubes compared to 67 genes upregulated in FSHD patient biopsies [20] and to 552 genes upregulated in *DUX4* expressing myotubes over non-expressing mytoubes [22].

**Figure S4.**
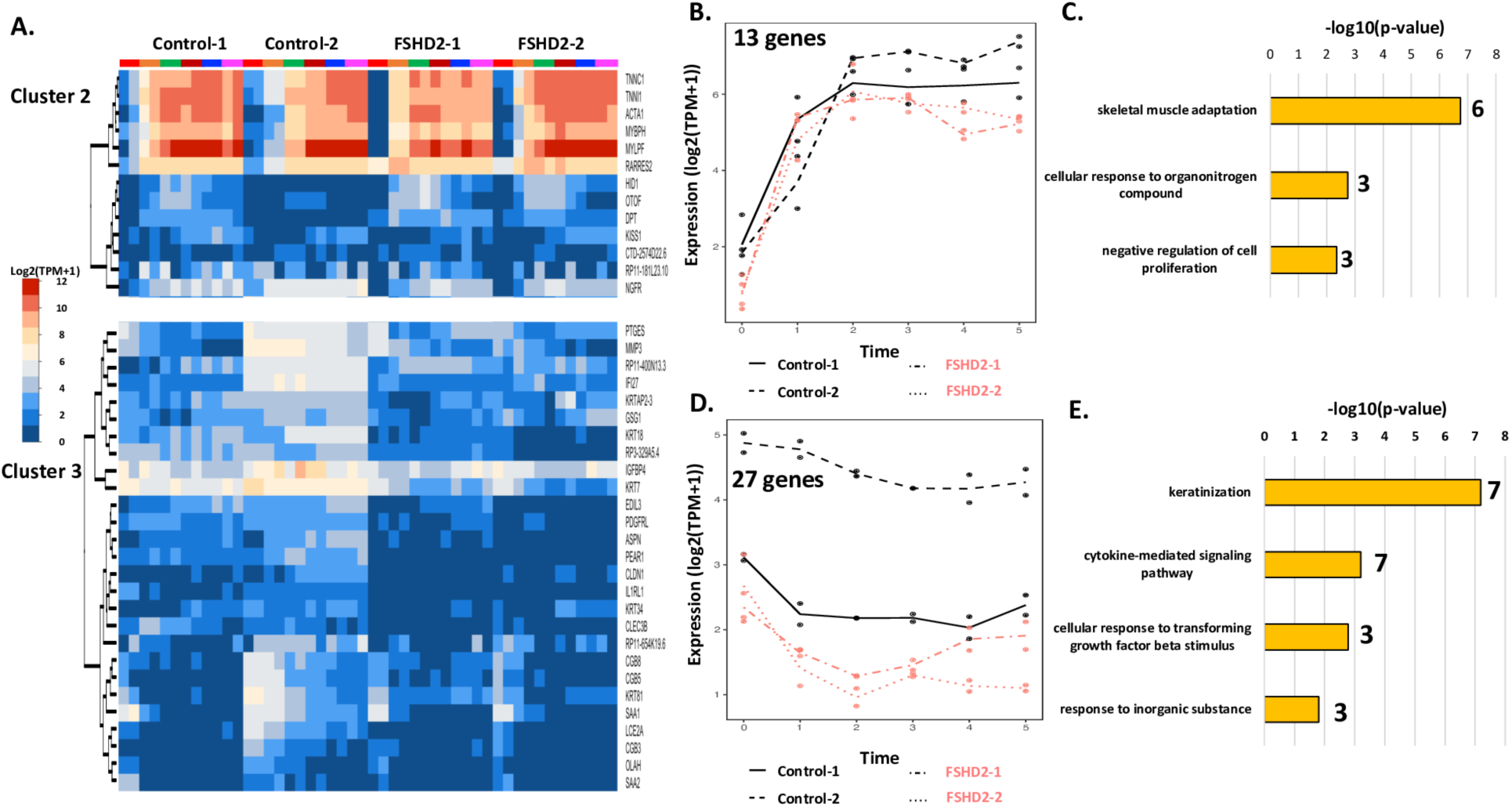
Differentially expressed genes clustered based on temporal expression differences (corresponds to Figure 2). **(A)** 103 differentially expressed genes clustered into four clusters by maSigPro based on expression patterns during control and FSHD2 differentiation time-course. Genes in cluster 2 are upregulated during differentiation in both control and FSHD2 while genes in cluster 3 are upregulated in Control-2 compared with others. **(B)** Expression profiles of the 13 genes in cluster 2 across four cell types during differentiation. **(C)** Gene ontology terms associated with the 13 genes in cluster 1. **(D)** Expression profiles of the 27 genes in cluster 3 across four cell types during differentiation. **(E)** Gene ontology terms associated with the 27 genes in cluster 4.

**Figure S5.**
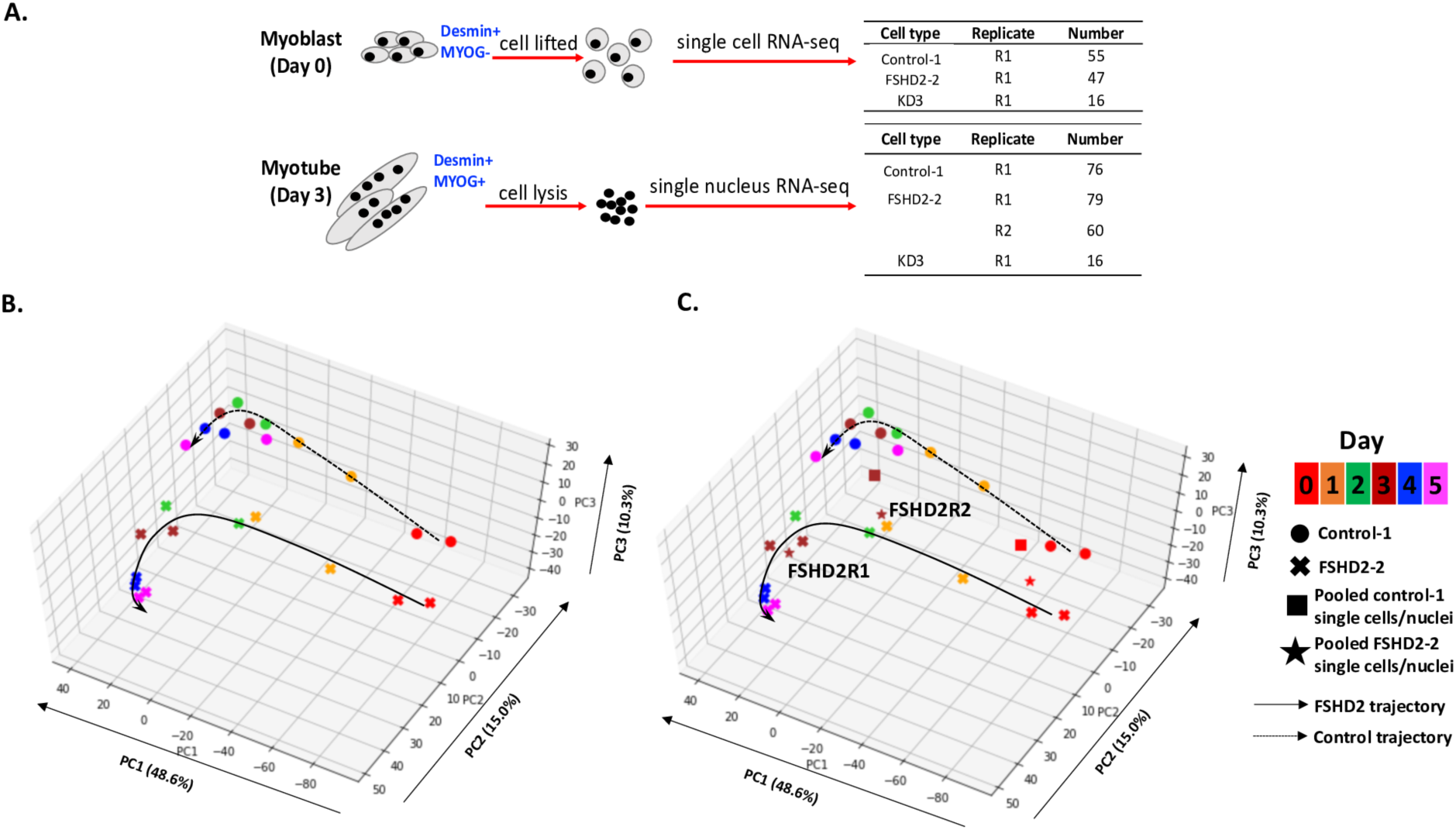
Overview of single-cell and single-nucleus samples and comparison with time-course. **(A)** Summary of single cells and single nuclei collected for sequencing. Single cells from myoblasts were selected to be *desmin*(+) *MYOG*(-) cells and retained for downstream analysis. Single nuclei from myotubes were selected to be *desmin*(+) *MYOG*(+) nuclei and retained for downstream analysis. **(B)** Principal component analysis (PCA) of Control-1 and FSHD2-2 myoblast differentiation time-course. Gene expression level was measured each day for duplicates by using RNA-seq. Cell types are labeled by shape, and time-points are labeled by color. **(C)** Incremental PCA on pooled Control-1 single cells and pooled FSHD2-2 single nuclei as well as bulk Control-1 and FSHD2-2 differentiation time-courses with the same dimensions as the PCA in (B).

**Figure S6.**
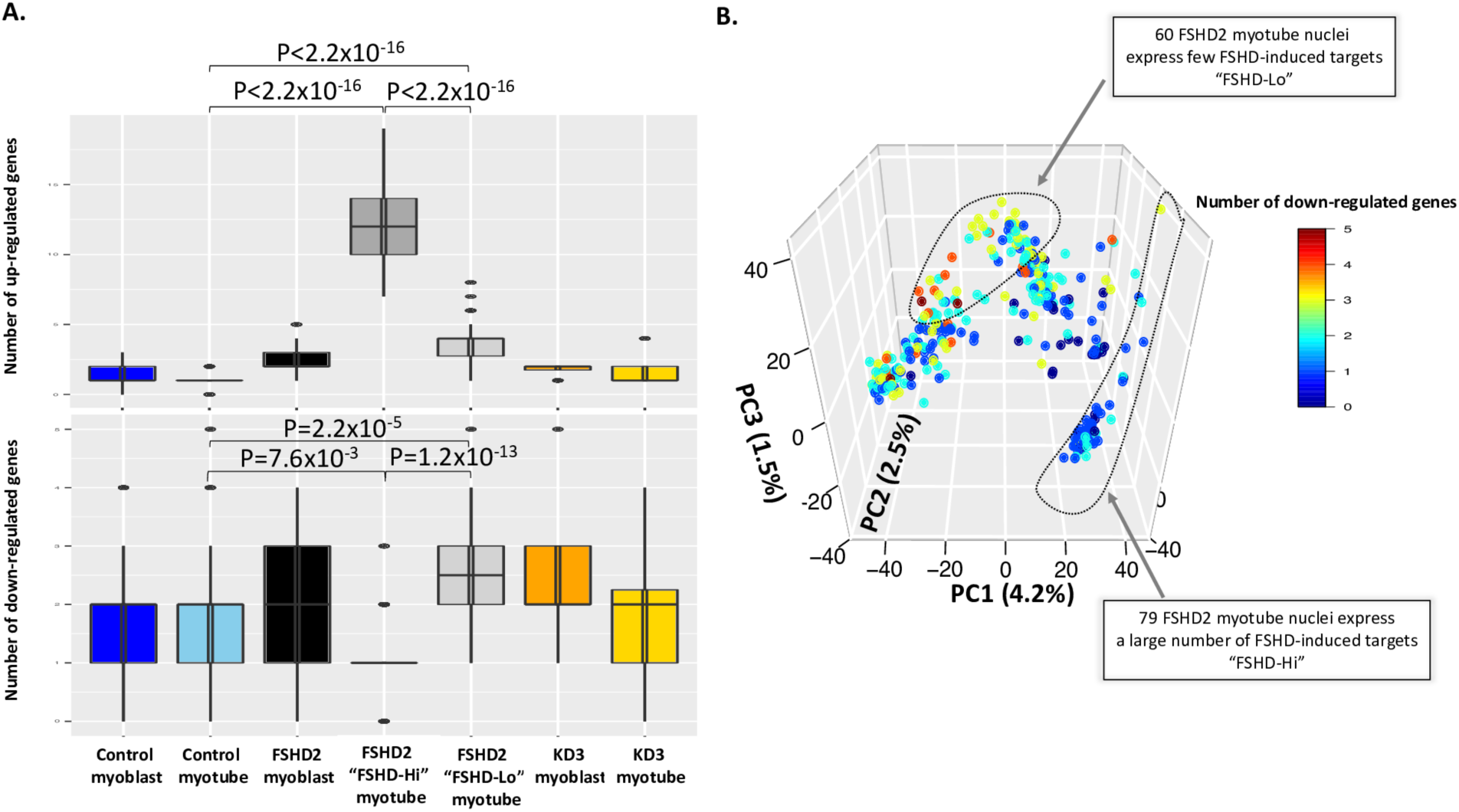
Differences in the number of up or downregulated genes from the time-course which are detected across sample types. **(A)** Comparison of the number of upregulated and downregulated genes in FSHD2 detected from time-course analysis across different cell types. **(B)** PCA of single-cell (for myoblast) and single-nucleus (for myotube) RNA-seq data for control, FSHD2 and KD3. Cells and nuclei are colored by the number of detected downregulated genes in FSHD2 determined from the time-course analysis.

**Figure S7.**
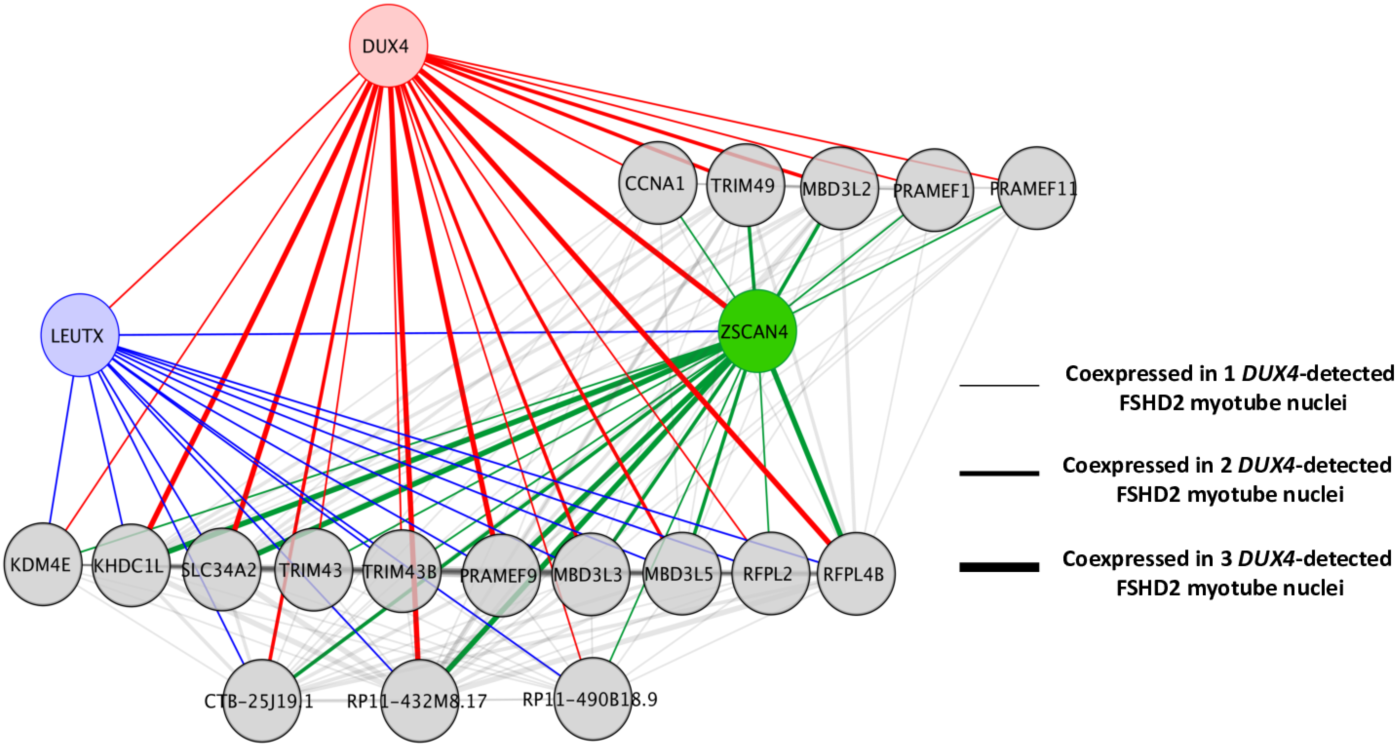
Coexpression network of genes in the three *DUX4*-detected nuclei. Twenty FSHD-induced genes are coexpressed with *DUX4*, two of which are transcription factors, *LEUTX* and *ZSCAN4*.

**Figure S8.**
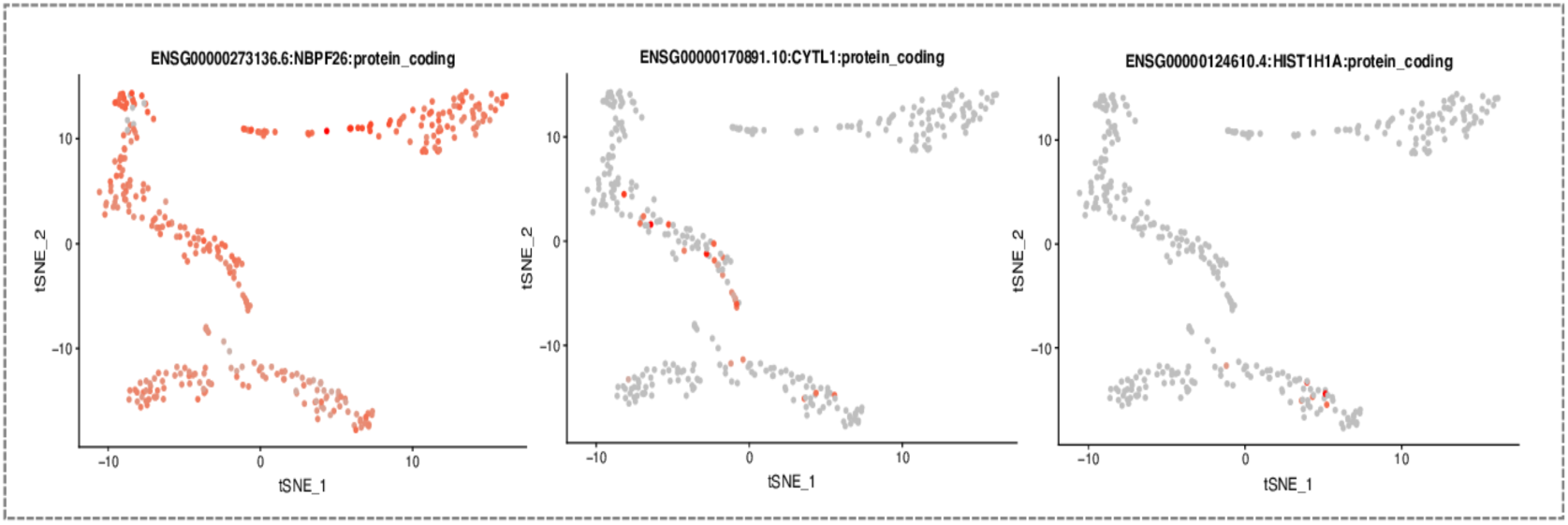
Expression of genes that start up-regulated at day 0 in FSHD2 time-course analysis across single cells and nuclei. Gene expression is visualized in t-SNE plots (same as Figure 4A) and cells/nuclei with expression of specific genes are colored in red.

**Figure S9.**
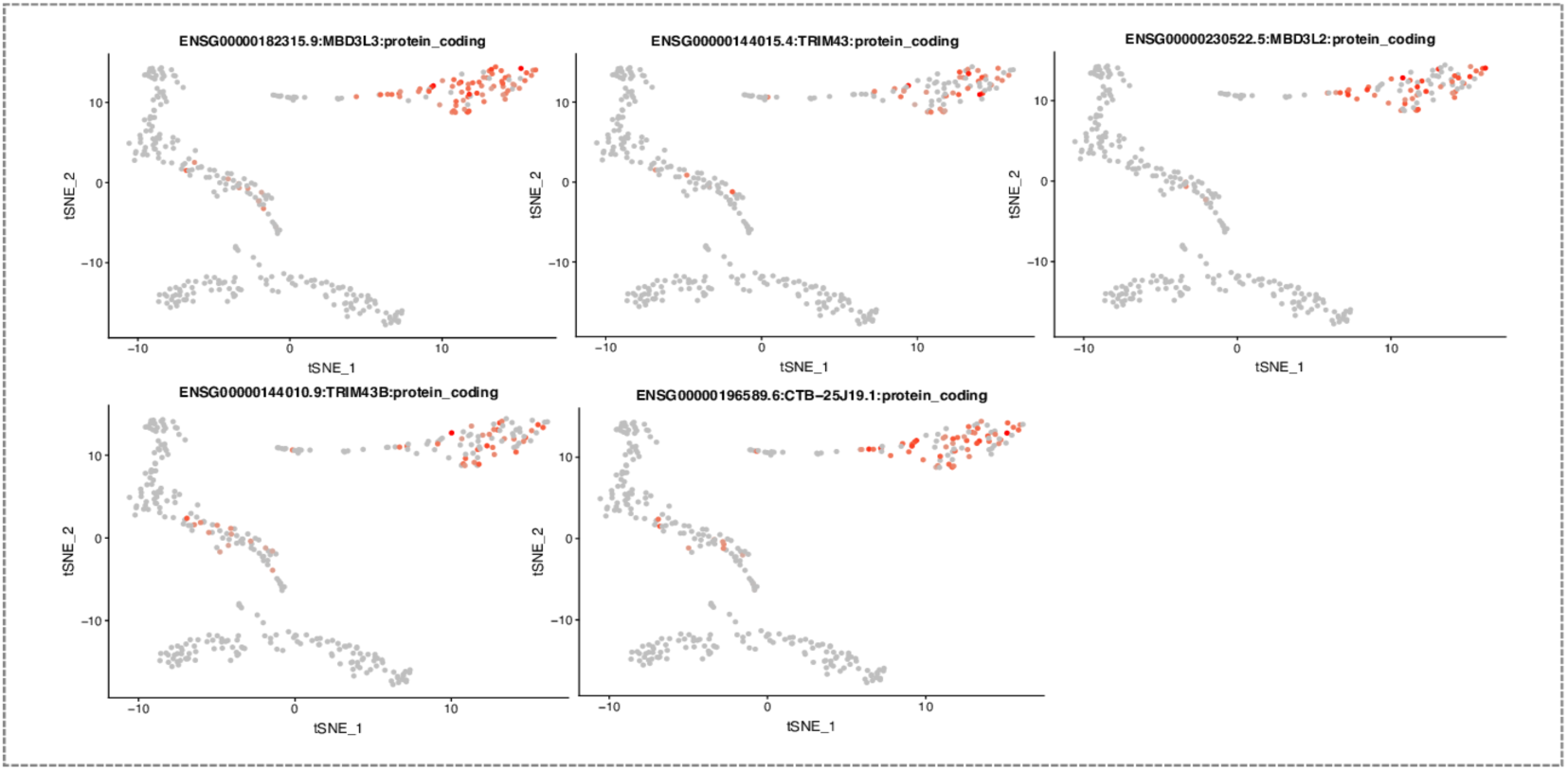
Expression of genes that start up-regulated at day 2 in FSHD2 time-course analysis across single cells and nuclei. Gene expression is visualized in t-SNE plots (same as Figure 4A) and cells/nuclei with expression of specific genes are colored in red.

**Figure S10.**
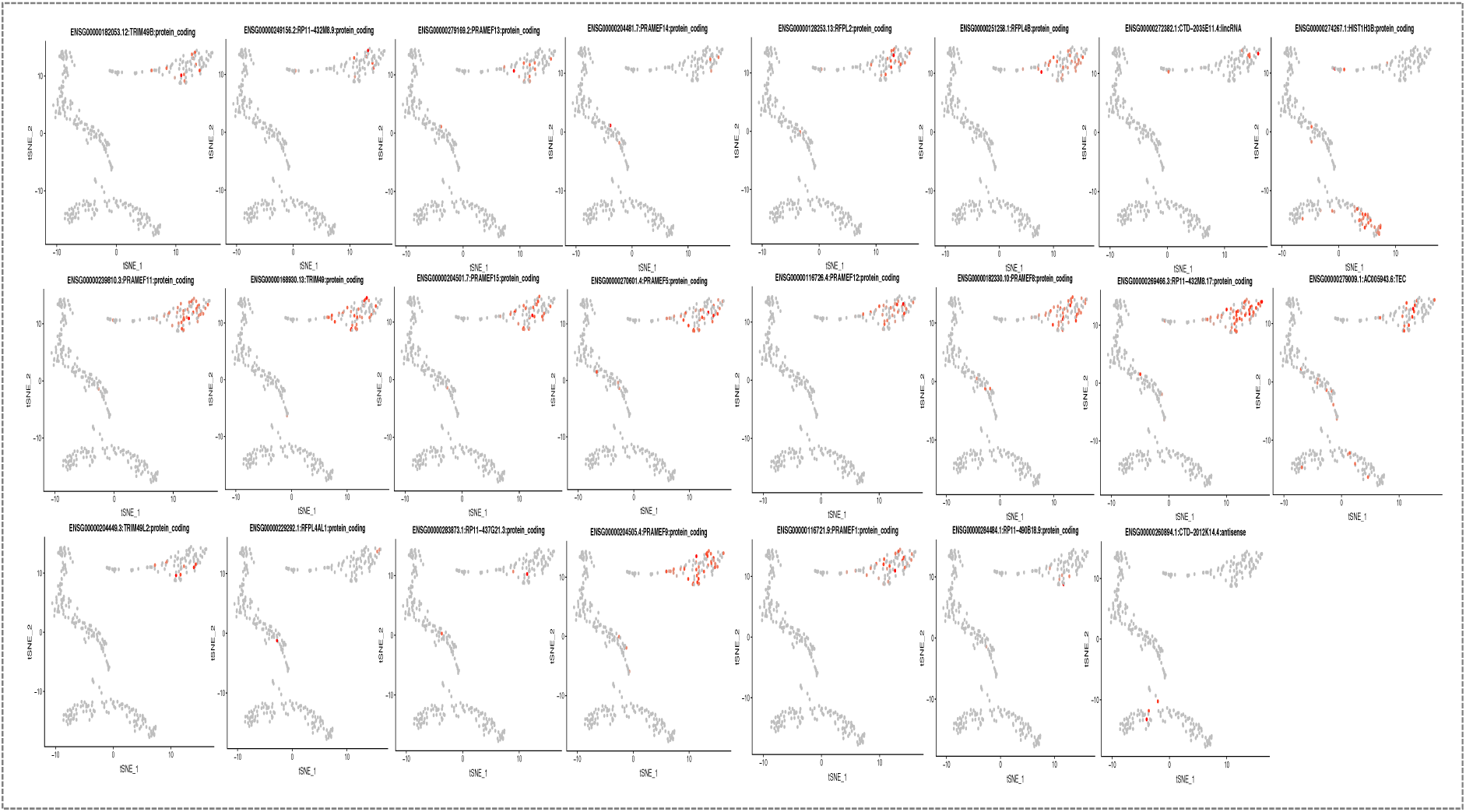
Expression of genes that start up-regulated at day 3 in FSHD2 time-course analysis across single cells and nuclei. Gene expression is visualized in t-SNE plots (same as Figure 4A) and cells/nuclei with expression of specific genes are colored in red.

**Figure S11.**
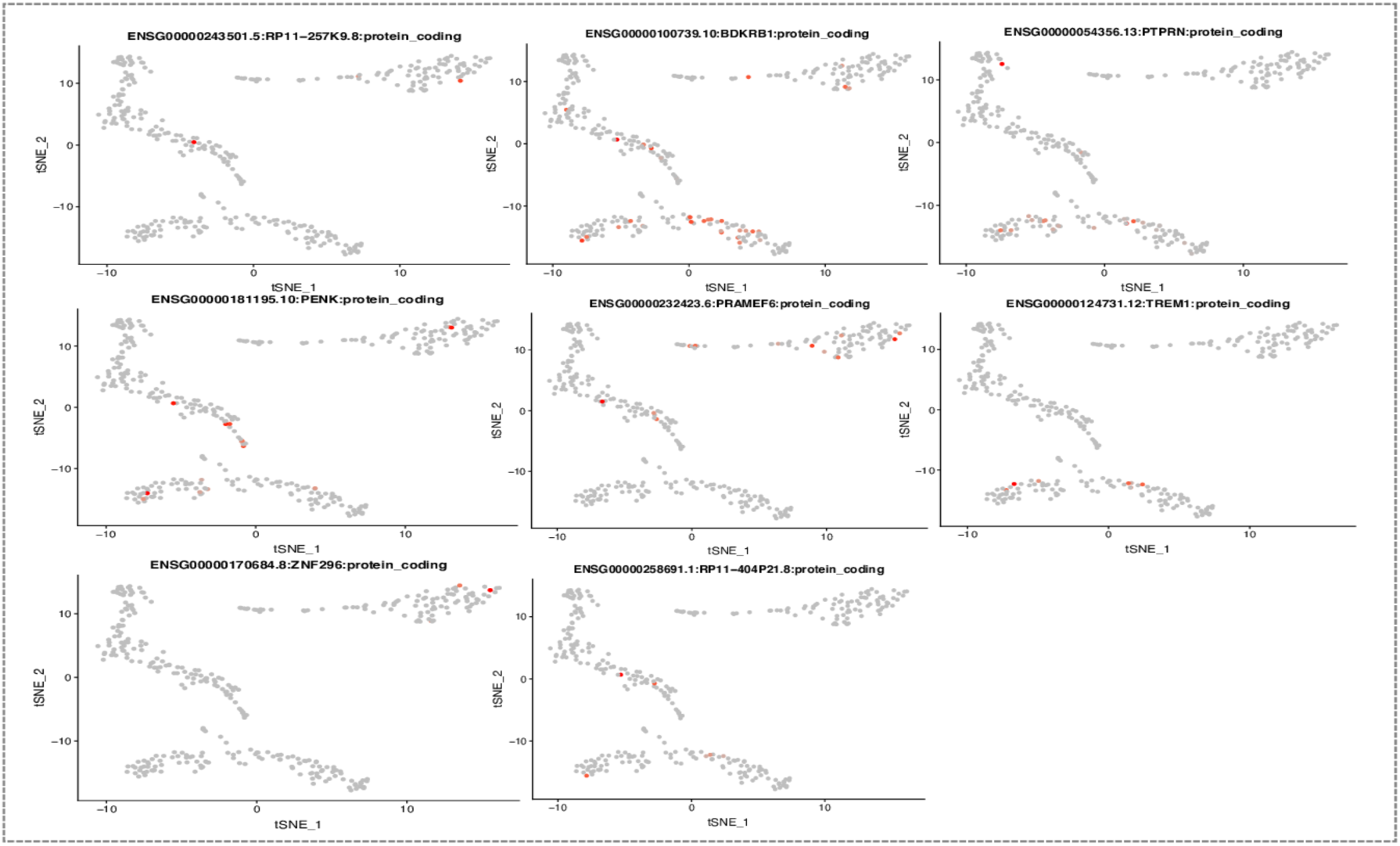
Expression of genes that start up-regulated at day 4 in FSHD2 time-course analysis across single cells and nuclei. Gene expression is visualized in t-SNE plots (same as Figure 4A) and cells/nuclei with expression of specific genes are colored in red.

**Figure S12.**
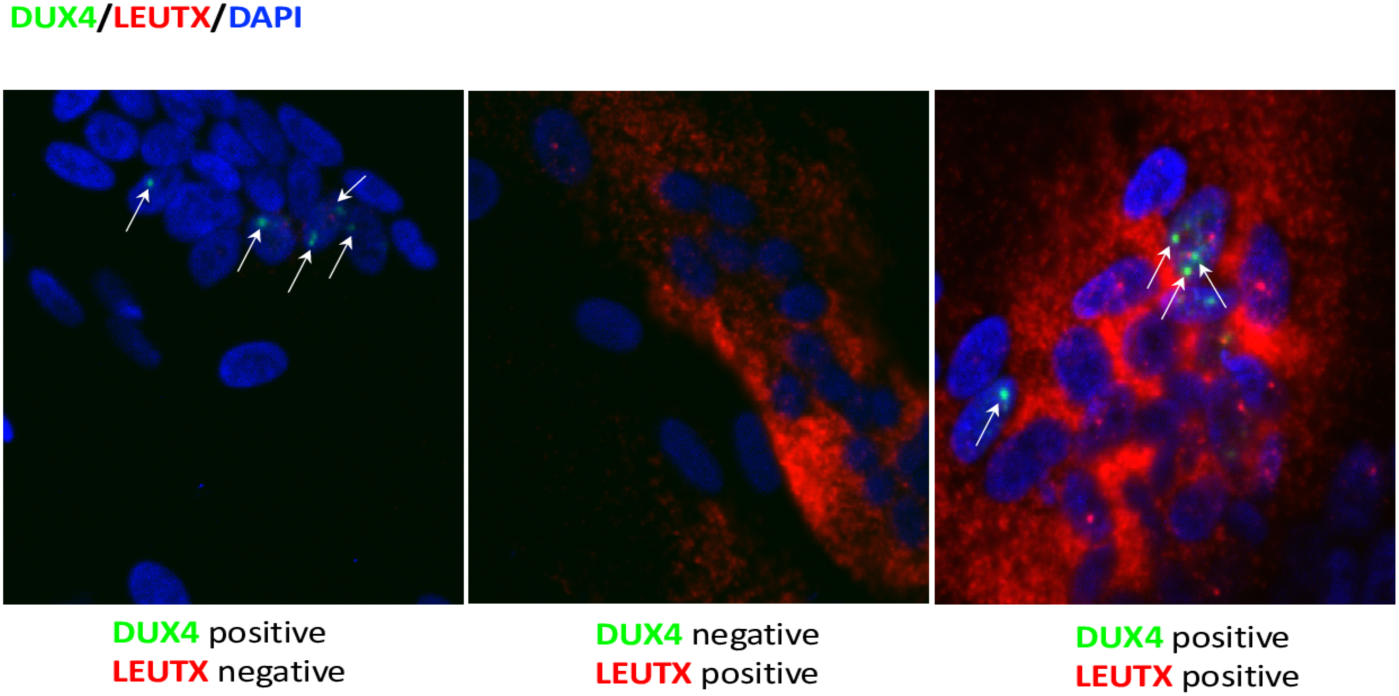
RNA FISH of *DUX4* and *LEUTX* in FSHD2 myotubes at day 3 of differentiation. *DUX4*, green; *LEUTX*, red; DAPI, blue.

**Figure S13.**
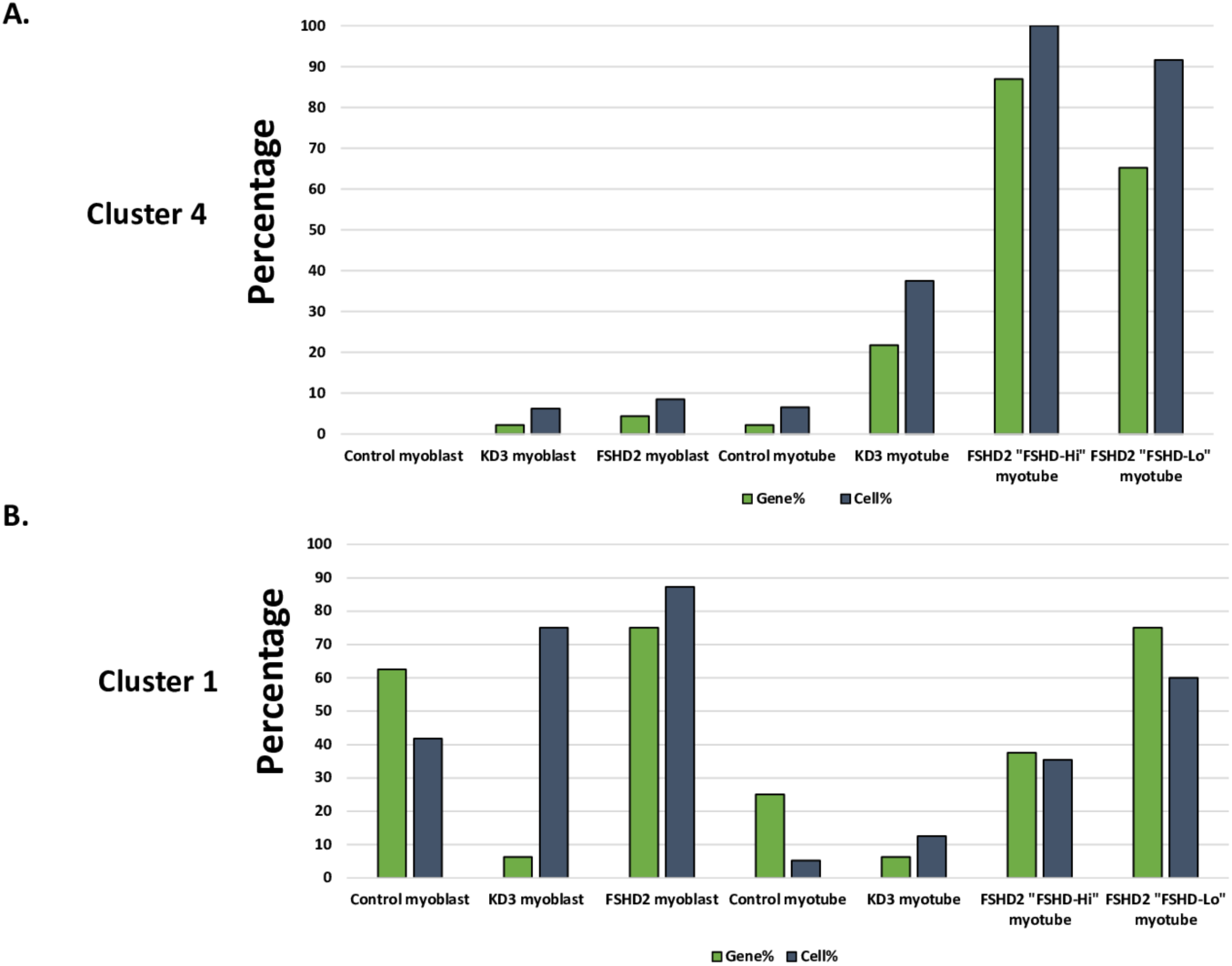
Histogram of early- and late-induced genes expressed in single cells and nuclei. **(A)** Percentage of single cells/nuclei (shown in dark gray) expressing late-induced genes that are identified in time-course analysis from clusters 4 and percentage of these genes being expressed (shown in green) across KD3, control and FSHD2. **(B)** Percentage of single cells/nuclei (shown in dark gray) expressing early-induced genes that are identified in time-course analysis from clusters 1 and percentage of these genes being expressed (shown in green) across KD3, control and FSHD2.

**Figure S14.**
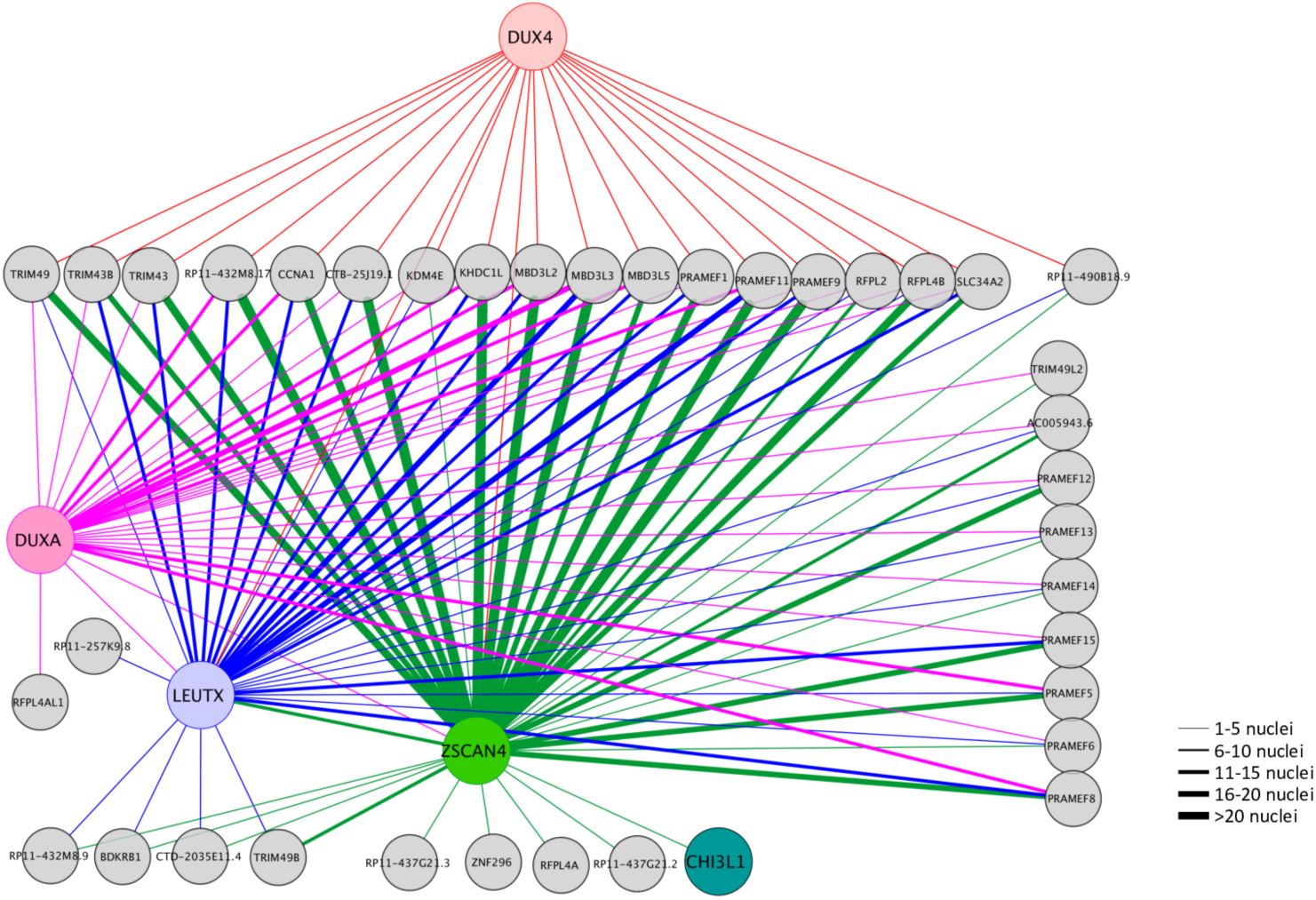
Coexpression network of FSHD-induced genes in “FSHD-Hi” myotube nuclei. FSHD-induced genes are coexpressed with *DUX4, DUXA, LEUTX* and *ZSCAN4* in “FSHD-Hi” nuclei.

**Figure S15.**
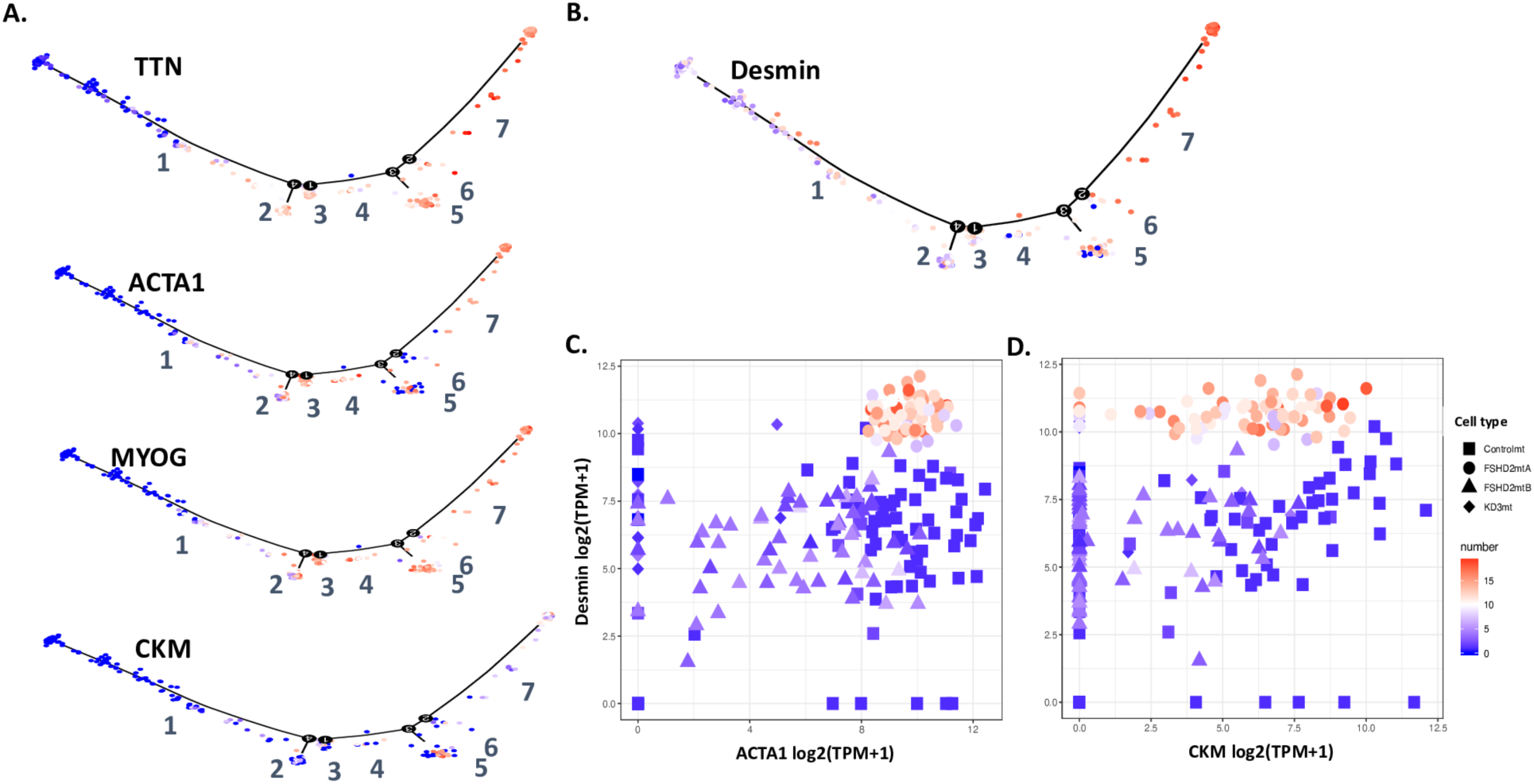
Myogenic gene expression changes in pseudotemporal ordering of 349 single cells/nuclei. Cells and nuclei are colored by the expression level of **(A)** myogenic markers and **(B)** *desmin*. Blue, low expression level; red, high expression level. **(C)** Scatter plot of gene expression between *ACTA1* and *desmin* in myotube nuclei. **(D)** Scatter plot of gene expression between *CKM* and *desmin* in myotube nuclei.

**Figure S16.**
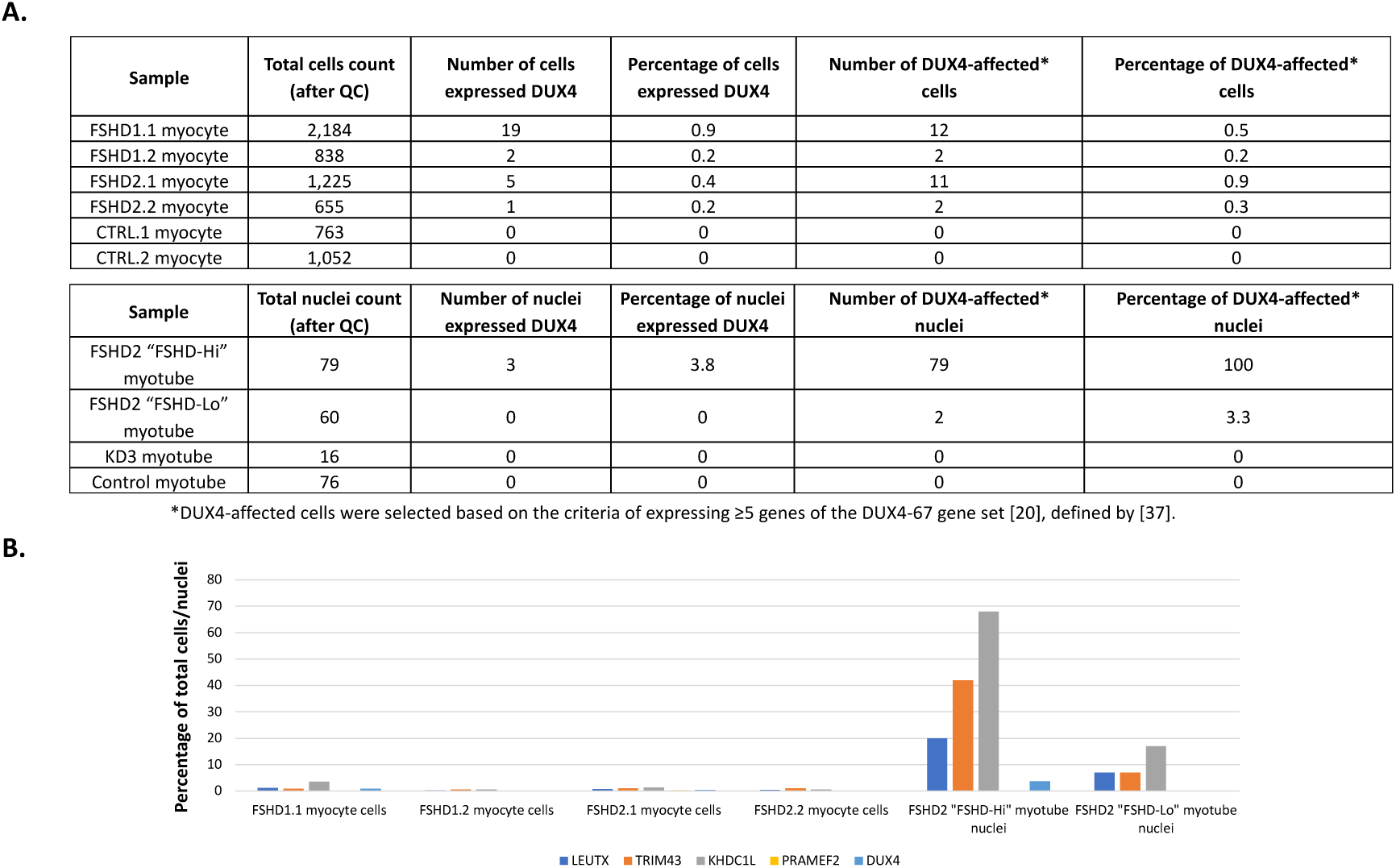
Comparison between published single cell FSHD myocyte RNA-seq data [37] and single nucleus FSHD myotube RNA-seq data in this study. **(A)** Number and percentage of *DUX4* expressing and affected myocyte single cells in published study [37] and myotube single nuclei in this study. **(B)** Percentage of total cells/nuclei expressing *DUX4* and 4 FSHD markers in myocyte single cells [37] and myotube single nuclei. 4 FHSD markers were selected from the published study [37] as a quality check.

Table S1. Principal components and distributed weights for 10,767 genes in Figure 1B.

Table S2. Gene expression level of 103 genes that are differentially expressed between FSHD2 and control during myoblast differentiation.

**Table S3.**
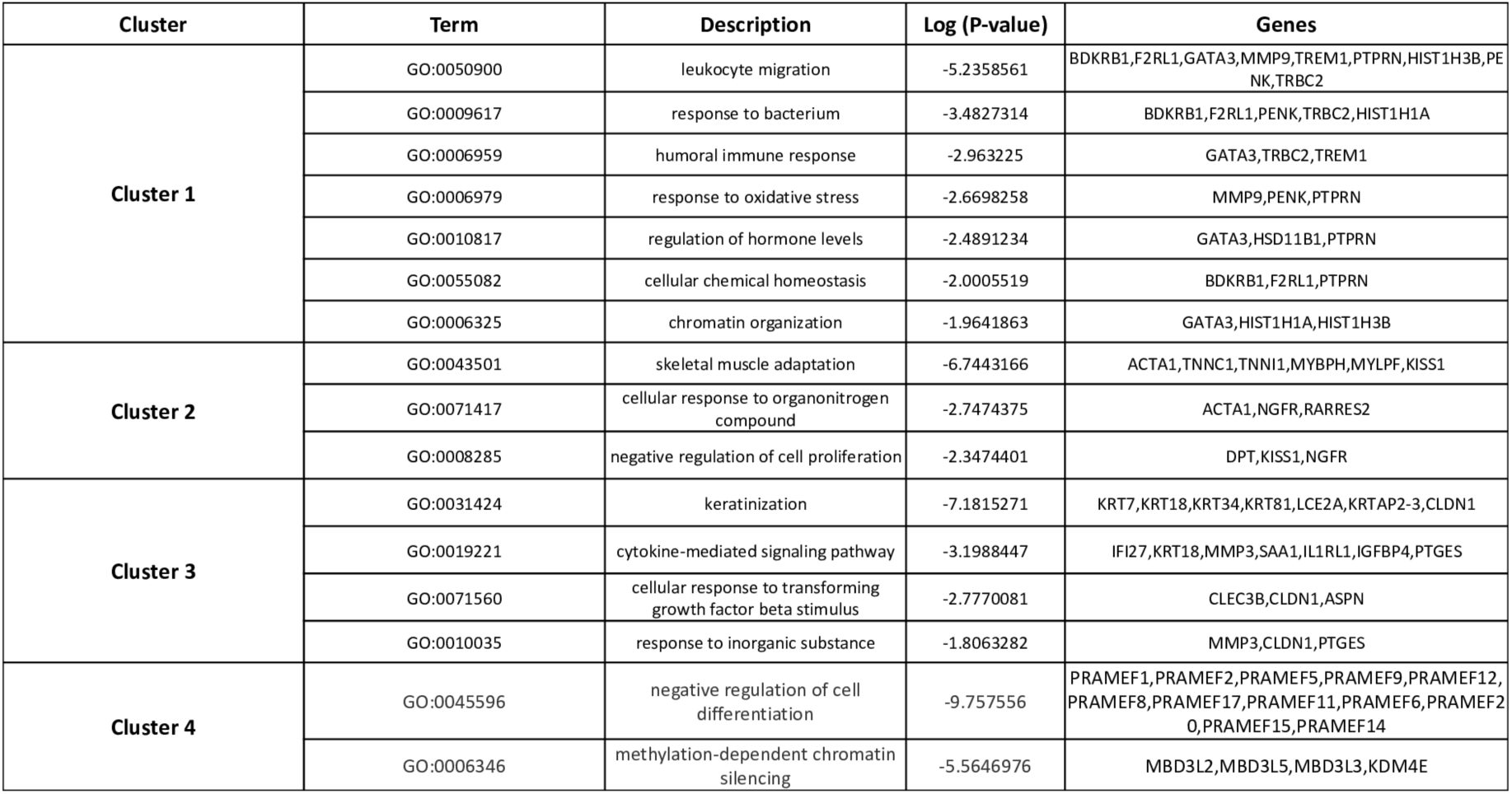
Gene ontology terms and enriched genes for the 4 clusters in Figures 2 and S4.

